# Multi-omic long-read sequencing reveals nucleosome control of transcriptional fidelity at the isoform level

**DOI:** 10.64898/2026.10.08.757737

**Authors:** Gali Bai, Namrita Dhillon, Brett Meissner, Colette Felton, Harrison D. Heath, Hinrich Boeger, Angela N. Brooks

**Affiliations:** Department of Biomolecular Engineering, University of California, Santa Cruz, Santa Cruz, California, 95064, United States of America; Department of Molecular, Cell, and Developmental Biology, University of California, Santa Cruz, Santa Cruz, California, 95064, United States of America

**Keywords:** Oxford Nanopore sequencing, single-molecule chromatin states, full-length isoform sequencing, nascent RNA long-read sequencing, co-transcriptional RNA processing, chromatin remodeling complexes

## Abstract

Transcription and RNA processing are tightly coupled to chromatin organization, yet how nucleosome positioning is coordinated across the promoter and gene body to shape alternative isoform expression remains poorly understood. This question has been difficult to address because short-read chromatin profiling cannot simultaneously capture promoter and gene-body chromatin states on the same DNA molecule, and short-read RNA sequencing cannot resolve full-length transcripts without requiring computational assembly. To address this, we deleted two conserved chromatin remodelers, *ISW1* and *CHD1,* in *Saccharomyces cerevisiae* and profiled chromatin fibers, nascent RNA, and full-length mature RNA using Oxford Nanopore long-read sequencing. We developed EpiFLAIR, a computational framework that assigns discrete long-range chromatin states across entire genes and integrates these states with isoform-level transcriptional outputs. Using EpiFLAIR, we show that single-molecule chromatin states together with sequence motifs are strong predictors of alternative TSS isoform expression.

Loss of *ISW1* and *CHD1* produced divergent transcriptional changes by activating transcriptionally silenced promoters while repressing active ones. Single-molecule chromatin profiling suggested that this redistribution of transcriptional activity arose, in part, from an anticorrelation between gene-body and promoter chromatin accessibility along the same chromatin fiber, a finding uniquely enabled by long-read single-molecule resolution. Furthermore, disruption of regular nucleosome spacing impaired RNAPII processivity and co-transcriptional splicing, consequently altering mature RNA levels and splicing isoform composition. Together, our findings demonstrate that nucleosomes maintain transcriptional fidelity during initiation, elongation, and splicing to regulate isoform expression. More broadly, long-read single-molecule chromatin profiling represents a powerful approach for linking chromatin organization to isoform-level gene regulation.

## INTRODUCTION

Transcription and RNA processing are tightly coordinated with chromatin organization (Li et al. 2007). The nucleosome, the basic device of DNA coiling, regulates DNA accessibility through its assembly, repositioning, and eviction. Active gene promoters are typically characterized by a nucleosome-depleted region (NDR) flanked by well-positioned −1 and +1 nucleosomes, with the +1 nucleosome facilitating pre-initiation complex assembly and transcription start site (TSS) selection (Lieleg et al. 2015; Wang et al. 2023; Baldi et al. 2020). Within coding regions, nucleosomes are regularly spaced to preserve chromatin integrity during transcription and promote efficient RNA polymerase II (RNAPII) progression (Uzun et al. 2021; Crickard et al. 2017; Kujirai and Kurumizaka 2020). Nucleosome occupancy is also higher over exons than introns, a pattern proposed to influence co-transcriptional splicing by modulating RNAPII elongation speed or recruiting splicing factors (Chen et al. 2010; Schwartz et al. 2009; Wu et al. 2025). Despite these proposed associations, how specific chromatin states shape RNA isoform diversity remains poorly understood.

This question has been technically intractable for two reasons. First, short-read chromatin profiling yields population-averaged nucleosome occupancy signals that cannot reveal whether accessibility at the promoter and accessibility within the gene body co-occur on the same DNA molecule (Liu and Conesa 2025). Short-read methods struggle to distinguish a mixed population with half open-promoter/closed-gene-body and half the reverse from one that is uniformly intermediate, even though these configurations carry profoundly different transcriptional implications. Second, short-read RNA sequencing cannot directly resolve full-length isoform structure without computational assembly, making it difficult to assign specific TSS choices, splicing patterns, and 3′ end positions to the same transcript molecule (Pardo-Palacios et al. 2024; Steijger et al. 2013). Together, these limitations have made it challenging to test whether distinct chromatin configurations on individual DNA molecules are causally linked to distinct isoform outcomes.

ATP-dependent chromatin remodelers provide an ideal experimental model to address this question. These remodelers use the energy of ATP hydrolysis to slide nucleosomes along DNA, establishing and maintaining uniform internucleosomal spacing that is essential both for protecting gene bodies from aberrant transcription initiation and for sustaining efficient RNAPII elongation (Clapier et al. 2017; Becker and Hörz 2002). Under a non-equilibrium model of chromatin organization, remodeler activity not only positions nucleosomes passively, but actively drives chromatin away from the sequence-determined thermodynamic equilibrium, amplifying differences in accessibility between active and inactive loci and suppressing cryptic initiation while sustaining transcriptional bursting at canonical promoters (Shelansky et al. 2024). This framework predicts that remodeler loss should shift nucleosome positioning toward a sequence-dependent equilibrium state and produce coordinated changes in isoform usage that are associated with the underlying chromatin configurations.

In *Saccharomyces cerevisiae*, Isw1 and Chd1 are the principal remodelers responsible for the intragenic nucleosome spacing during active transcription (Smolle et al. 2012; Ocampo et al. 2016). Isw1 acts preferentially within coding regions to dampen cryptic transcription (Tirosh et al. 2010), while Chd1 cooperates with the transcription machinery and histone chaperones to reassemble nucleosomes in the wake of RNAPII passage (Hennig et al. 2012). Loss of *ISW1* and *CHD1* disrupts gene-body nucleosome spacing and increases cryptic transcription (Ocampo et al. 2016; Smolle et al. 2012; Hennig et al. 2012). However, previous studies only measured population-averaged chromatin alterations, and no studies have yet integrated that with long-read RNA-seq to study the association between long-range chromatin states and specific isoform outcomes.

To characterize genome-wide single-molecule chromatin states and their association with RNA isoforms, we deleted *ISW1* and *CHD1* in *S. cerevisiae* and used Oxford Nanopore long-read sequencing to profile chromatin fibers, nascent RNA, and mature RNA simultaneously. We developed EpiFLAIR, a computational framework that classifies individual chromatin fibers into discrete states based on coordinated promoter and gene-body accessibility, and links these states to isoform-level gene expression. Our results reveal that chromatin remodelers actively maintain the transcriptional landscape by coordinating nucleosome organization across promoters and gene bodies, and that disruption of this coordination reshapes isoform usage genome-wide.

## RESULTS

### Chromatin remodeler loss compresses the transcriptome’s expression range

To study the consequences of remodeler loss on transcription and chromatin structure, we deleted two major ATP-dependent chromatin remodelers, *ISW1* and *CHD1* (Methods). We profiled chromatin accessibility (2 replicates), nascent RNA (2 replicates), and full-length mature RNA (3 replicates) in both wild-type and *isw1*Δ *chd1*Δ double-mutant cells using long-read nanopore sequencing (Figure 1A). In addition, short-read Illumina sequencing of total RNA (3 replicates) was performed in wild-type and mutant cells to quantify changes in gene expression (Figure 1A). Differential expression analyses of short-read total RNA-seq data identified a modest number of genes that were altered in the mutant, with 508 genes up-regulated while 179 genes down-regulated (Supplementary Figure 1C, Supplementary Figure 1D).

**Figure 1.**
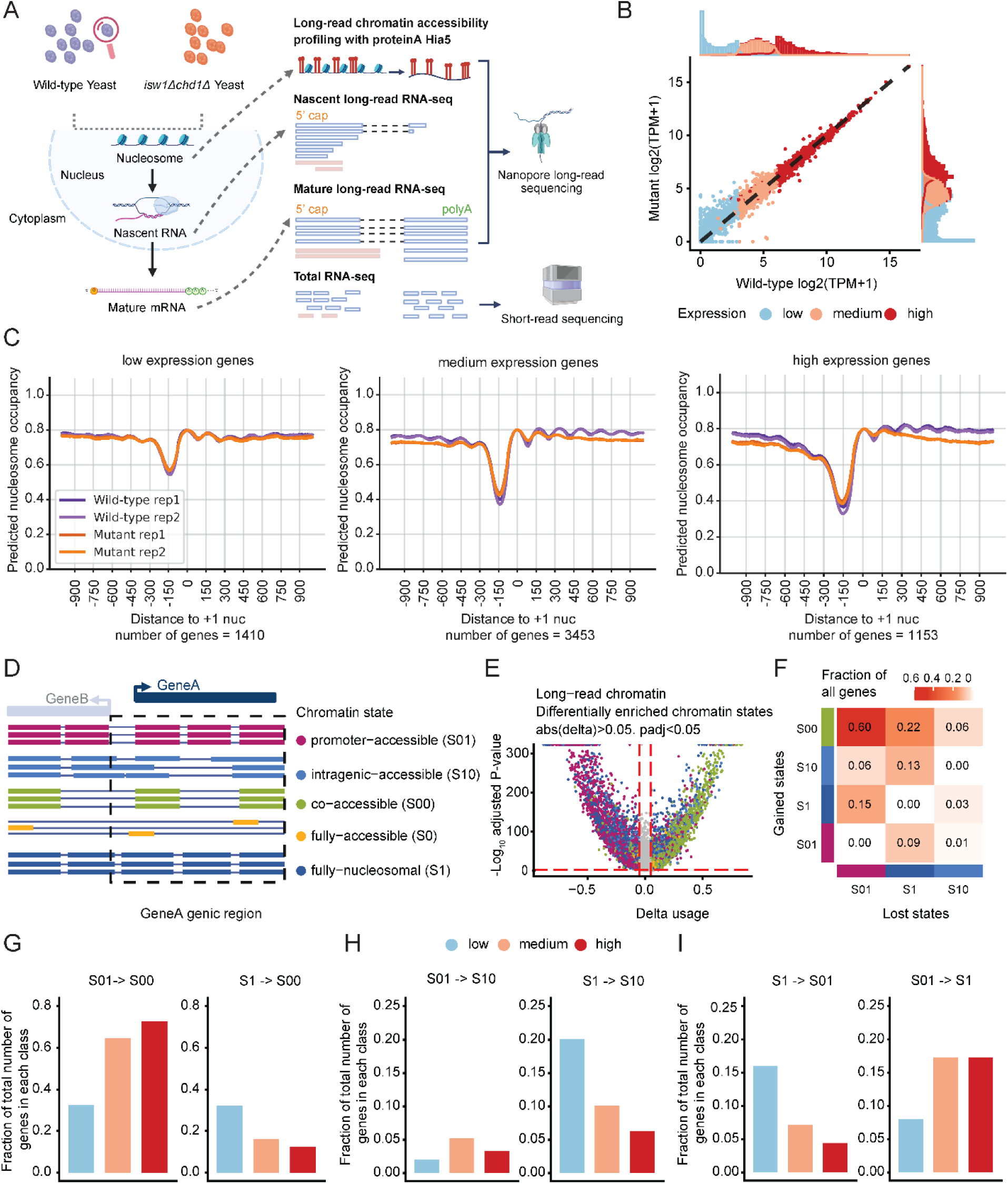
Experimental design and single-molecule long-range chromatin state analysis in *isw1*Δ *chd1*Δ mutant yeast. (A) Schematic of the multi-omic experimental design. Wild-type and *isw1*Δ *chd1*Δ double mutant *S. cerevisiae* were profiled by long-read chromatin accessibility sequencing (pA-Hia5 methyltransferase footprinting with Oxford Nanopore sequencing), long-read nascent RNA sequencing, full-length mature RNA sequencing, and short-read total RNA sequencing. Created in BioRender. Bai, G. (2026) https://BioRender.com/6o32n5d. (B) Scatter plot and histogram comparing gene-level expression (log_2_ TPM) between mutant and wild-type cells. Each point represents one gene, colored by baseline expression class (low, medium, high). (C) Aggregate nucleosome occupancy profiles centered on the +1 nucleosome dyad for low-, medium-, and high-expression genes in wild-type and *isw1*Δ *chd1*Δ mutant cells. Profiles are averaged across all reads per expression class. (D) Schematic of the five single-molecule chromatin state classifications defined by EpiFLAIR based on the coordinated nucleosome occupancy across the promoter and gene body on the same chromatin fiber: promoter-accessible (S01), in which only the promoter is open, intragenic-accessible (S10), in which only the gene body is open, co-accessible (S00), in which both promoter and gene body are simultaneously open, fully accessible (S0), in which more than 90% of the genic region is nucleosome-free, and fully nucleosomal (S1), in which both promoter and gene body are closed. Created in BioRender. Bai, G. (2026) https://BioRender.com/grw4z0j. (E) Volcano plot of differentially enriched chromatin states between *isw1*Δ *chd1*Δ mutant and wild-type cells. Each point represents a chromatin state at a given gene. The x-axis shows the delta usage difference in chromatin state fraction between mutant and wild-type, and the y-axis shows the −log_10_ adjusted p-value. Points above the significance threshold (|delta| > 0.05, padj < 0.05) are colored by chromatin state identity. (F) Transition matrix showing the proportion of genes that shift from one chromatin state in wild-type to another in the *isw1*Δ *chd1*Δ mutant. Cell color intensity reflects the fraction of genes undergoing each transition. (G-I) Bar plots showing the fraction of genes transitioning between specific chromatin state pairs in the mutant relative to wild-type, stratified by baseline expression class. Each bar represents the fraction of genes in that expression class undergoing the indicated state transition.

To further parse these expression changes, we stratified genes into low, medium, and high expression groups based on wild-type expression levels (Figure 1B, Methods). Genes with low basal expression were preferentially up-regulated in the mutant, whereas a small fraction of highly expressed genes tended to be down-regulated, resulting in slightly reduced global expression variance (Figure 1B, Supplementary Figure 1E, Supplementary Figure 1F). Specifically, 34.9% of low-expression genes were up-regulated, compared with 4.7% of medium- and 1.4% of high-expression genes. Conversely, 0.4% of low-expression genes were down-regulated, compared with 3.0% of medium- and 6.0% of high-expression genes (Supplementary Figure 1E). Together, these results demonstrate that loss of *ISW1* and *CHD1* compresses the range of transcriptional output by differentially impacting genes based on their basal expression level.

### Chromatin states recapitulated the baseline transcriptional landscape

To investigate how chromatin organization relates to transcriptional activity, we sequenced chromatin from 2 wild-type replicates using long-read nanopore sequencing with pA-Hia5 methylation of accessible adenosines and inferred nucleosome occupancy on individual molecules using fibertools (Jha et al. 2024) (Supplementary Figure 2A). Sequencing yielded approximately 3M mapped reads per sample, with an average coverage of 300x, a median read length of ∼1.5 kb, and read quality of Q25 (Supplementary Figure 2B, Supplementary Table 1). To process and integrate the long-read chromatin data with long-read RNA-seq, we developed a computational method, EpiFLAIR (Methods).

Using EpiFLAIR, we identified nucleosome-depleted regions (NDRs) on individual fibers and determined the position of the +1 nucleosome for each gene (Methods, Supplementary Figure 2C). To assess whether our nucleosome predictions suggest associations between chromatin states and baseline gene expression, we compared nucleosome occupancy profiles across low-, medium-, and high-expression genes in wild-type cells. Aligning chromatin fibers at the +1 nucleosome of each gene and averaging occupancy across all genes, we observed the expected ∼150 bp nucleosome footprint and characteristic periodicity of regularly spaced nucleosomes (Supplementary Figure 2D). This periodicity gradually decayed into the gene body beyond the well-positioned +1, +2, and +3 nucleosomes, as expected from the statistical positioning of nucleosomes (Kornberg and Stryer 1988). When we sorted profiles by expression class, NDR depth was prominent in high- and medium-expression genes but markedly reduced in low-expression genes (Supplementary Figure 2E). Interestingly, individual chromatin fibers of the same gene displayed a wide range of promoter accessibilities, with the fraction of fiber configurations reflecting the gene’s transcriptional state, consistent with single-molecule EM analyses of promoter nucleosome dynamics (Supplementary Figure 2F, Supplementary Figure 2G, Supplementary Figure 2H) (Brown et al. 2013; Wolff et al. 2021). These results confirm that our long-read chromatin profiling captures the expected relationship between nucleosome organization and gene expression at both population and single-molecule resolution, and highlight the importance of classifying individual chromatin fibers into discrete states rather than relying on population-averaged signals.

### Perturbation of nucleosome phasing and chromatin accessibility was transcription-dependent

We next compared aggregate wild-type and mutant chromatin profiles. In contrast to the regularly spaced nucleosome profiles in wild-type cells, mutant samples showed substantial disruption of gene-body nucleosome spacing as previously shown (Supplementary Figure 3A) (Xu et al. 2025; Ocampo et al. 2016). Interestingly, the data also suggested that decreased gene-body nucleosome occupancy coincided with increased nucleosome occupancy at promoters, potentially reflecting redistribution of nucleosomes from gene bodies into promoter-proximal regions (Supplementary Figure 3A).

When stratified by expression group, gene-body nucleosome periodicity in low-expression genes showed minimal change upon remodeler loss, but was highly pronounced in highly expressed genes (Figure 1C, Supplementary Figure 3B). Notably, highly transcribed genes with the most prominent NDRs showed the most pronounced gains in promoter nucleosome occupancy, accompanied by a concomitant loss of translational positioning of intragenic nucleosomes (Supplementary Figure 3C). These findings indicate that chromatin reorganization caused by *ISW1* and *CHD1* loss is correlated with the transcriptional activity of that locus, consistent with the idea that these remodelers are preferentially recruited to, and therefore most consequential at, highly transcribed loci (Gkikopoulos et al. 2011).

### Single-molecule chromatin profiling reveals coordinated promoter and gene body nucleosome redistribution upon remodeler loss

Long-read chromatin profiling enables reconstruction of nucleosome configurations along individual DNA fibers, allowing structural correlations between promoter and intragenic regions to be observed on the same molecule. Leveraging this capability, we investigated whether accessibility changes at promoters and within gene bodies are coordinated at the single-molecule level. We applied EpiFLAIR to classify individual chromatin fibers into five discrete states based on the coordinated nucleosome occupancy across the promoter and gene body (Methods, Figure 1D, Supplementary Figure 3D). Four states reflect the combination of promoter and gene body accessibility: promoter-accessible (S01), in which the promoter is open but intragenic regions are closed; intragenic-accessible (S10), in which the promoter is closed but at least one exon carries an accessible region; co-accessible (S00), in which both promoter and intragenic regions are simultaneously open on the same fiber; and fully nucleosomal (S1), in which both regions are closed. The fifth state, fully accessible (S0), is distinguished from co-accessible (S00) to define fibres in which more than 90% of the entire genic region is accessible, representing a global loss of nucleosome occupancy rather than the discrete dual-opening characteristic of S00. Comparing the fraction of chromatin state S01 between wild-type and mutant cells, we found high reproducibility within replicates (Pearson r = 0.78 for wild-type; r = 0.75 for mutant) but lower correlation between conditions (Pearson r ≤ 0.65) (Supplementary Figure 3E).

Differential usage analyses of chromatin states showed that the mutant has a global increase in the abundance of the co-accessible state (S00) and a corresponding reduction in the promoter-accessible state (S01) (Figure 1E, Supplementary Figure 4A, Supplementary Table 2), with 60% of all genes shifting from S01 to S00 upon loss of *ISW1* and *CHD1* (Figure 1F). When stratified by baseline expression level, the magnitude of this S01 to S00 transition scaled with baseline transcriptional activity (Figure 1G, Supplementary Figure 4B), consistent with preferential remodeler recruitment to highly transcribed loci and proportionately greater disruption upon their loss.

Beyond this dominant transition, ∼19% of genes showed an increase in gene body accessibility, accompanied by a decrease in accessibility over the promoter (S10) on the same chromatin fiber (Figure 1F). The increased abundance of the S10 state was enriched in low- and medium-expression genes rather than highly expressed ones (Figure 1H, Supplementary Figure 4B), suggesting that intragenic accessibility can arise independently of promoter state in low- and medium-expression genes. In contrast to the previous chromatin state transition, the fully nucleosomal state (S1) showed opposing behavior by expression class: it was depleted in low-expression genes, where remodeler loss de-represses previously silenced loci, but increased in medium- and high-expression genes, where nucleosome redistribution from gene bodies toward promoter-proximal regions may occlude accessibility (Figure 1F, Figure 1I, Supplementary Figure 4B).

Notably, the overall fraction of methylated reads and total nucleosome occupancy were minimally changed between wild-type and mutant at single-molecule resolution (Supplementary Figure 4C), indicating that *ISW1* and *CHD1* loss drives genome-wide nucleosome redistribution rather than net eviction. These expression-dependent, coordinated changes in promoter and intragenic chromatin states are uniquely observable via single-molecule long-read profiling, whereas population-averaged short-read data would obscure the co-occurrence of these states on individual fibers.

### Long-read sequencing of nascent and mature RNA resolves isoform-level transcriptional dynamics

Prior studies examining the relationship between chromatin accessibility and transcriptional output have largely operated at the gene level, reporting modest or inconsistent correlations between nucleosome occupancy changes and differential gene expression in remodeler mutants (Gkikopoulos et al. 2011). We reasoned that this disconnect may reflect a fundamental limitation of gene-level analysis, where chromatin perturbations that shift isoform usage without changing much of total RNA would be invisible to gene-level quantification.

To directly test whether chromatin state changes alter RNA processing and isoform expression, we profiled nascent RNA and full-length mature RNA by nanopore long-read sequencing in wild-type and double mutant cells (Figure 1A). Full-length mature RNA profiling using the TeloPrime kit selectively enriches transcripts that are both 5′-capped and polyadenylated (Figure 2A). For nascent RNA, because the library preparation does not include a dedicated 5′-cap enrichment step, we developed a post-alignment filtering strategy to identify and retain only reads from capped transcripts. Inspection of aligned nascent RNA reads revealed that a fraction of reads carried non-genome-encoded 5′ guanosine bases, which were reported to define the 5 ′ end of cDNA (Supplementary Figure 5A, Supplementary Table 3) (Ohtake et al. 2004). This signature was further confirmed by comparison with matched public CAGE-seq data, establishing that reads with additional 5′ Gs represent capped nascent transcripts with known transcription initiation sites (Supplementary Figure 5B). Reads carrying poly(A) tails, indicative of mature RNA, were also discarded from the nascent RNA analyses (Supplementary Figure 5A, Supplementary Table 3). Together, this filtering approach allowed us to distinguish bona fide nascent transcripts from contaminating mature RNA, and to assign each nascent read to a precise transcription start site (Figure 2B, Methods).

**Figure 2.**
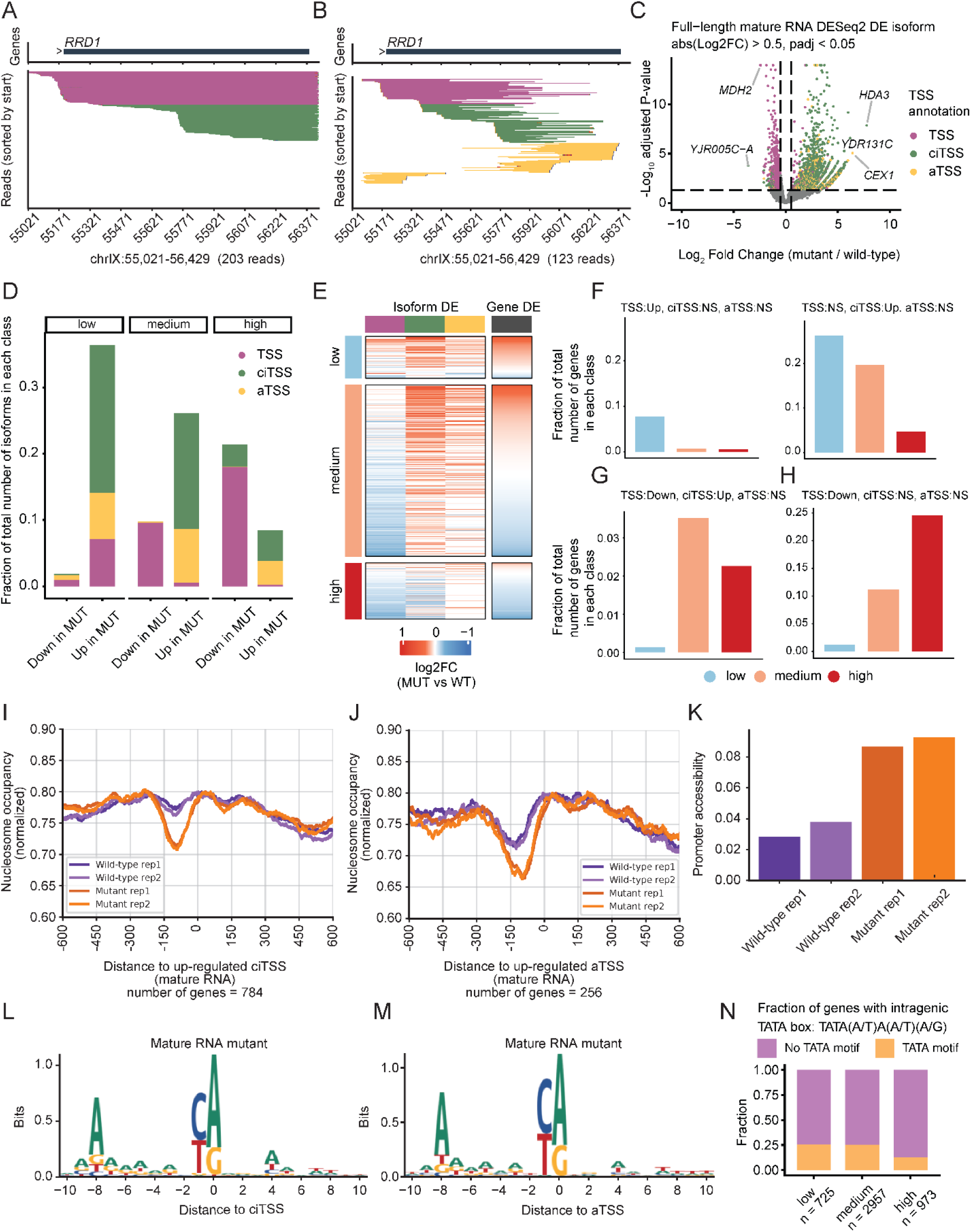
Loss of *ISW1* and *CHD1* induces cryptic intragenic and antisense transcription initiation from promoter-like sequences. (A-B) Representative genome browser track views of full-length mature RNA (A) and nascent RNA (B) reads aligned to the *RRD1* locus in *isw1*Δ *chd1*Δ mutant cells. Individual reads are colored by TSS class: TSS (pink), cryptic intragenic TSS (ciTSS, green), and antisense TSS (aTSS, yellow). Gene annotation is shown on each track with exons as boxes and the direction of transcription indicated by arrows. (C) Volcano plot showing differentially expressed TSS isoforms between the *isw1*Δ *chd1*Δ mutant and wild-type cells from mature RNA. Each point represents one TSS isoform. Points above the significance threshold (|log_2_FC| > 0.5, padj < 0.05) are colored by TSS annotation. (D) Bar plots showing the fraction of differentially expressed TSS isoforms from mature RNA in each baseline expression class, stratified by direction of change (up- or down-regulated in the mutant). (E) Heatmap of isoform-level and gene-level log_2_ fold change (mutant vs wild-type) across expression classes. Each row represents one gene, with columns showing the fold change for the TSS, ciTSS, and aTSS isoforms, alongside the total gene-level expression change, illustrating how isoform switching contributes to changes in total gene output. (F-H) Bar plots showing the fraction of genes with each combination of co-occurring TSS isoform changes from mature RNA, stratified by baseline expression class. For each gene, the expression change of each TSS class (TSS, ciTSS, and aTSS) in the *isw1*Δ *chd1*Δ mutant relative to wild-type is classified as down-regulated (Down), up-regulated (Up), or not significantly changed (NS). Genes are grouped by their combination of outcomes across all three TSS classes. (I-J) Aggregate nucleosome occupancy profiles centered at up-regulated ciTSSs (I) and aTSSs (J) in the *isw1*Δ *chd1*Δ mutant. (K) Quantification of chromatin accessibility at the nucleosome-depleted region centered at up-regulated ciTSSs, compared across wild-type and mutant samples. (L-M) Sequence logos of the initiator (Inr) motif centered at ciTSSs (L) and aTSSs (M) in the mutant. The conserved PyPu dinucleotide at the −1/+1 position with an A at −8 recapitulates the canonical yeast TSS consensus. (N) Fraction of genes containing an intragenic TATA box motif (TATA(A/T)A(A/T)(A/G)), stratified by baseline expression class.

Nascent RNA reads had a median length of ∼369 bp, with approximately 1.7M reads per sample (Supplementary Figure 5C, Supplementary Table 3). Full-length mature RNA reads had a median length of ∼741 bp, with approximately 2.5M reads per sample (Supplementary Figure 5D and 5E, Supplementary Table 4). Given that the median gene length in yeast is 1044 bp (Supplementary Figure 1B), full-length coverage of the mature RNA library was biased toward shorter genes due to the limited processivity of the reverse transcriptase and short fragment cDNA amplification bias during library preparation. The nascent RNA library did not exhibit this bias due to its shorter transcript length (Supplementary Figure 5F, 5G). Consequently, genes longer than ∼3,000 bp were covered by less than one full-length mature RNA read on average in the Teloprime data (Supplementary Figure 5H and 5I).

We quantified isoform and gene expression from both mature and nascent RNA libraries using custom pipelines (Supplementary Figure 6A, Supplementary Figure 6B). Expression estimates were more highly correlated within sequencing methods than between them (Supplementary Figure 6C). In wild-type cells, nascent RNA expression correlated more strongly with short-read total RNA (Pearson r = 0.74) than did mature RNA (Pearson r = 0.73) due to the read-length bias in the mature RNA libraries (Supplementary Figure 6D, 6E). Despite some library-specific biases, comparing the relative enrichment of nascent and mature RNA populations provides a complementary view of transcriptional and post-transcriptional RNA processing dynamics.

### Loss of *ISW1* and *CHD1* induces cryptic intragenic and antisense transcription initiation

Previous studies have shown that loss of *ISW1* and *CHD1* induces cryptic intragenic and antisense transcription with microarray data (Smolle et al. 2012). To more precisely characterize changes in transcription initiation in *isw1*Δ *chd1*Δ mutant cells, we annotated three classes of transcription start site (TSS) isoforms for all non-intron-containing genes: TSSs (TSS) initiating from the annotated start site, cryptic intragenic TSSs (ciTSS), and antisense TSSs (aTSS) on the opposite strand (Figure 2A, 2B, Methods). TSS usage was quantified from both nascent and full-length mature RNA sequencing data (Methods). Differential expression analysis using DESeq2 revealed a global shift in transcriptional initiation in mutant cells: ciTSS and aTSS isoforms were more up-regulated, while TSS isoform expression decreased in both full-length and nascent RNA (Figure 2C, Supplementary Figure 7A, Supplementary Table 5 and 6). By profiling both 5′-capped nascent and full-length mature RNA, our study captures precise cryptic transcription initiation sites from both short-lived transcripts and those that successfully proceed through elongation and processing to yield stable polyadenylated isoforms.

Stratifying these changes by expression class, low-expression genes showed the greatest up-regulation in the mutant in the full-length RNA-seq, with 36.4%, 26.1%, and 8.5% of low, medium, and high expression genes, respectively, being up-regulated (Figure 2D). Down-regulation was most pronounced in highly expressed genes: 1.8%, 9.8%, and 21.4% in the low, medium, and high expression groups, respectively. Similar trends were also observed in nascent RNA (Supplementary Figure 7B). These results are consistent with the differential expression patterns observed in the short-read RNA-seq (Supplementary Figure 1E). To understand how TSS isoform-level changes contribute to total gene expression, we examined co-occurring TSS changes within individual genes in the full-length and nascent RNA-seq data. In low-expression genes, transcription increased broadly across all TSS classes, collectively driving higher total expression (Figure 2E, 2F, Supplementary Figure 7C-E). In medium-expression genes, decreases in TSS isoforms were frequently offset by increases in ciTSS or aTSS isoforms, resulting in relatively stable total expression but pronounced isoform switching (Figure 2E, 2G, Supplementary Figure 7C-E). In high-expression genes, both TSS and ciTSS isoforms decreased within the same genes, producing an overall reduction in transcriptional output (Figure 2E, 2H, Supplementary Figure 7C-E).

Together, our data suggest that loss of *ISW1* and *CHD1* reduces differences in transcriptional output across TSSs. Combining this observation with our chromatin data that show redistribution of nucleosomes away from cryptic promoters and a modest closing of canonical promoters suggests a model in which total transcriptional output is preserved by activation of previously repressed initiation sites and dampening of transcription from the most active loci in wild-type cells.

### Cryptic intragenic transcription initiates from promoter-like sequences within gene bodies

We next asked whether the chromatin alterations observed in the mutant underlie cryptic intragenic transcription initiation. For each gene, we annotated ciTSSs and aTSSs in mutant cells using nascent and mature long-read RNA data, respectively, and examined aggregated nucleosome occupancy centered at these sites in up-regulated genes. The chromatin landscape at ciTSSs and aTSSs closely resembled that of canonical promoters: initiation sites were more likely to be nucleosome-depleted than flanking intragenic sequences, with well-positioned +1 and +2 nucleosomes immediately downstream (Figure 2I, 2J, Supplementary Figure 8A-B). Consistent with this interpretation, these sites were more accessible in mutant than in wild-type cells (Figure 2K), indicating that remodelers normally suppress cryptic initiation by maintaining nucleosome occupancy over the gene body.

To determine whether ciTSSs and aTSSs share promoter sequence features with TSSs, we performed motif analysis across all three classes. All three TSSs were enriched for the initiator (Inr) motif (PyPu at −1/+1) with an A at the −8 position, consistent with the previously described yeast TSS consensus (Figure 2L, 2M, Supplementary Figure 8C-F) (Lu and Lin 2019; Zhu et al. 2024; Qiu et al. 2020; Wei et al. 2019; Doris et al. 2018). Interestingly, despite having the strongest gene-body nucleosome disruption in the mutant, highly expressed genes showed the lowest levels of ciTSS activity despite loss of nucleosomes across the gene body (Figure 1C, Figure 2D, Supplementary Figure 7B). We hypothesized that gene bodies of highly transcribed genes are intrinsically depleted of promoter-like sequences, providing a sequence-encoded safeguard against aberrant initiation in regions subject to frequent nucleosome disassembly during active transcription. Examination of core promoter motif enrichment, including TATA box (TATA(A/T)A(A/T)(A/G)) (Basehoar et al. 2004) and Inr elements, across expression-stratified genes revealed that high-expression genes had the lowest fraction of TATA motifs within their gene bodies, whereas Inr elements are quite ubiquitously distributed (Figure 2N, Supplementary Figure 8G). This pattern further supports that sequence composition and nucleosome occupancy act together to protect highly transcribed gene bodies from cryptic initiation.

### Chromatin structure perturbations quantitatively account for changes in TSS isoform expression

To determine whether chromatin state changes are associated with corresponding shifts in mature RNA isoform composition, we computed Spearman correlations between alterations in chromatin state (Δ fraction) and full-length isoform expression changes (log2FC) across genes (Methods). The expected biological relationships were recovered: TSS expression correlated positively with the promoter-accessible state (S01) and negatively with the fully nucleosomal state (S1), while ciTSS expression correlated positively with the intragenic-accessible (S10) and co-accessible (S00) states (Supplementary Figure 9A, 9B). These relationships suggest that the abundance of chromatin states is quantitatively correlated with isoform expression.

Stratifying this chromatin-isoform coupling by baseline expression level revealed a clear dependence on transcriptional context. In low- and medium-expression genes, chromatin state changes were strongly concordant with isoform usage shifts: depletion of the promoter-accessible state (S01) and enrichment of intragenic-accessible (S10) and co-accessible (S00) states correlated with decreased TSS isoform usage and increased ciTSS and aTSS isoform usage (Figure 3A, 3E). In highly expressed genes, however, these correlations were much weaker (Figure 3I). As discussed above, this lack of correlation may be due to the paucity of promoter-like sequences in strongly expressed genes and the reported ability of strong upstream promoters to outcompete downstream cryptic sites and suppress aberrant initiation (Lee et al. 2021).

**Figure 3.**
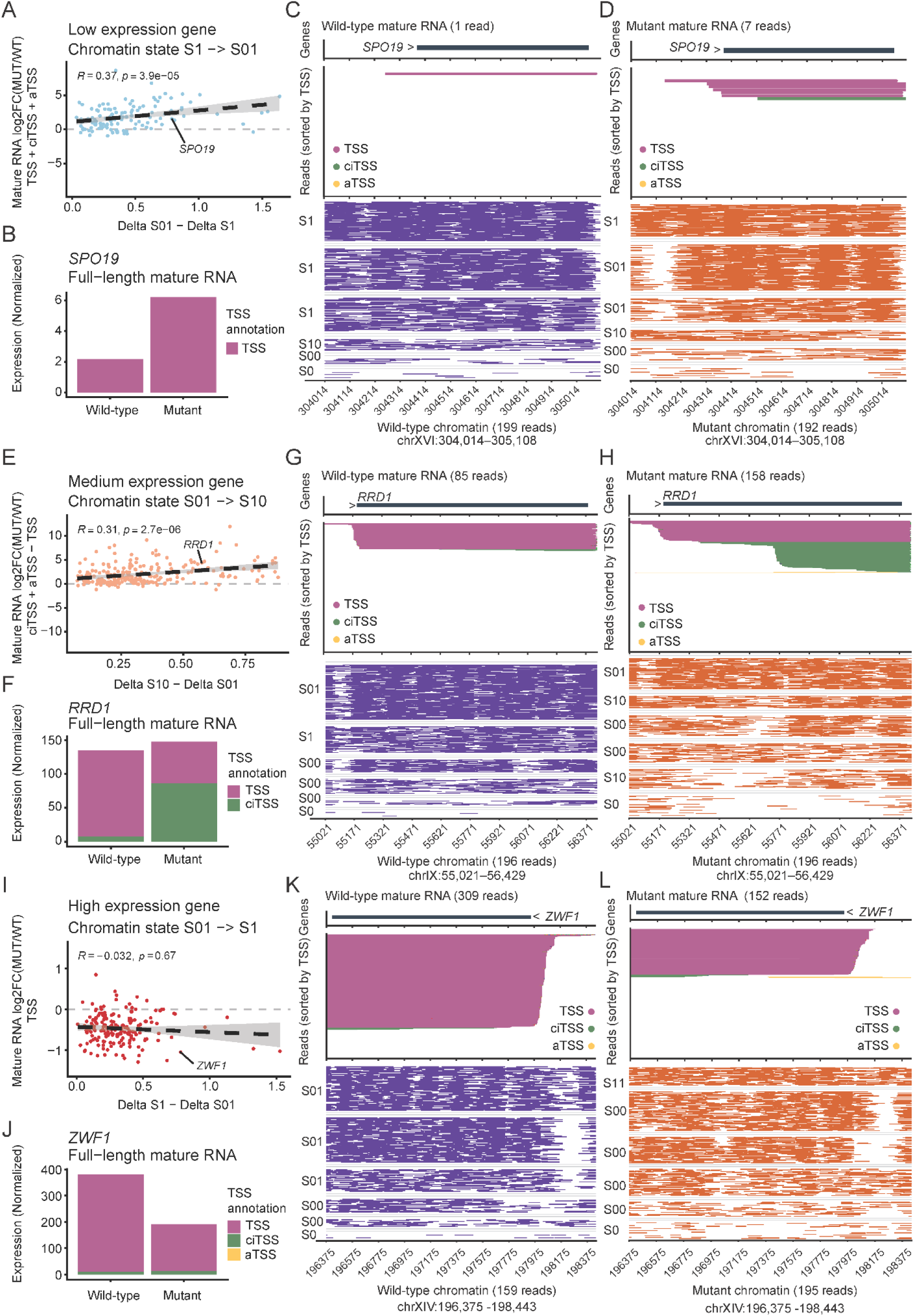
Single-molecule chromatin state changes correlate with full-length isoform expression in *isw1*Δ *chd1*Δ mutant yeast. (A) Scatter plot showing the Spearman correlation between the change in chromatin state fraction (Δ promoter-accessible S01 - fully-nucleosomal S1) and the log_2_ fold change of all TSS isoform expression (TSS + ciTSS + TSS) across low-expression genes. Each point represents one gene. (B) Bar plot of full-length mature RNA TSS isoform expression in the *SPO19* gene in wild-type and *isw1*Δ *chd1*Δ mutant cells. Expression values are variance-stabilizing transformed (VST) counts from DESeq2, shown separately for TSS, ciTSS, and aTSS isoforms. (C-D) Genome browser track views of full-length mature RNA reads (top) and single-molecule chromatin fibers (bottom) at the *SPO19* locus (chrXVI:304,014-305,108) in wild-type (C) and *isw1*Δ *chd1*Δ mutant (D) cells. Individual transcripts are colored by TSS annotation. (E) Scatter plot showing the Spearman correlation between the change in chromatin state fraction (Δ intragenic-accessible S10 - promoter-accessible S01) and the log_2_ fold change of ciTSS + aTSS - TSS full-length mature RNA isoforms across medium-expression genes. Each point represents one gene. (F) Bar plot of full-length mature RNA TSS isoform expression in the *RRD1* gene in wild-type and *isw1*Δ *chd1*Δ mutant cells. Expression values are variance-stabilizing transformed (VST) counts from DESeq2, shown separately for TSS, ciTSS, and aTSS isoforms. (G-H) Genome browser track views of full-length mature RNA reads (top) and single-molecule chromatin fiber maps (bottom) at the *RRD1* locus (chrIX:55,021-56,429) in wild-type (G) and *isw1*Δ *chd1*Δ mutant (H) cells. (I) Scatter plot showing the Spearman correlation between the change in chromatin state fraction (Delta fully-nucleosomal S1 - promoter-accessible S01) and the log_2_ fold change of TSS full-length mature RNA isoforms across high-expression genes. Each point represents one gene. (J) Bar plot of full-length mature RNA TSS isoform expression in the *MAP1* gene in wild-type and *isw1*Δ *chd1*Δ mutant cells. Expression values are variance-stabilizing transformed (VST) counts from DESeq2, shown separately for TSS, ciTSS, and aTSS isoforms. (K-L) Genome browser track views of full-length mature RNA reads (top) and single-molecule chromatin fiber maps (bottom) at the *ZWF1* gene locus (chrXIV:196,375-198,443) in wild-type (G) and *isw1*Δ *chd1*Δ mutant (H) cells.

We examined representative genes from each expression class to illustrate how these chromatin-isoform relationships manifest at individual loci. *SPO19* is a low-expression gene: In wild-type cells, nucleosomes block the entire genic region, including its promoter, effectively repressing transcription (Figure 3B and 3C). In the mutant, increased promoter accessibility coincides with a gain in transcription of the TSS isoform (Figure 3D). *RRD1* is an example of a medium-expression gene where remodeler loss produces isoform switching (Figure 3F). In wild-type cells, the *RRD1* promoter is accessible in the majority of chromatin fibers while the gene body remains largely nucleosomal, and transcription is dominated by TSS isoform expression (Figure 3G). In the mutant, nucleosomes redistribute from the gene body toward the promoter, activating intragenic NDRs while reducing promoter accessibility on the same fibers. This coordinated chromatin state transition, from promoter-accessible (S01) to intragenic-accessible (S10), is accompanied by a reciprocal shift in isoform usage, with decreased canonical TSS isoform expression and increased ciTSS isoform expression (Figure 3H), directly illustrating how single-molecule chromatin state changes predict isoform switching at this locus. Finally, the highly expressed gene *ZWF1* shows less obvious coupling of chromatin accessibility and transcription (Figure 3J-L). In the mutant, as gene-body chromatin becomes more accessible while the promoter becomes more closed, TSS isoform usage declines without any compensatory gain in ciTSS or aTSS transcription.

While chromatin accessibility changes are associated with shifts in initiation site choice in most low- and medium-expression genes, they do not account for the disproportionately large reductions of mature RNAs observed in highly expressed genes, which were down-regulated despite modest changes in promoter accessibility (Supplementary Figure 9C).

### Loss of nucleosome spacing impairs RNAPII processivity

To better understand why some highly expressed genes produce less mature RNA in the mutant even when their promoters remain accessible, we compared isoform expression changes between mutant and wild-type cells across nascent and full-length mature RNA libraries. Although changes in isoform expression were well correlated between the two libraries (Pearson r = 0.6), ∼38% of TSS isoforms showed discordant regulation: up-regulated in nascent RNA yet down-regulated in full-length mature RNA in mutant cells (Figure 4A, 4B). Discordantly regulated isoforms were enriched among highly expressed genes, the same set of genes that showed the greatest disruption of gene-body nucleosome spacing, pointing to a processivity defect as the underlying cause (Supplementary Figure 10A). We also observed that the discrepancy between nascent and mature RNA was significantly less pronounced for ciTSS and aTSS isoforms (Supplementary Figure 10B), consistent with their lower baseline expression levels.

**Figure 4.**
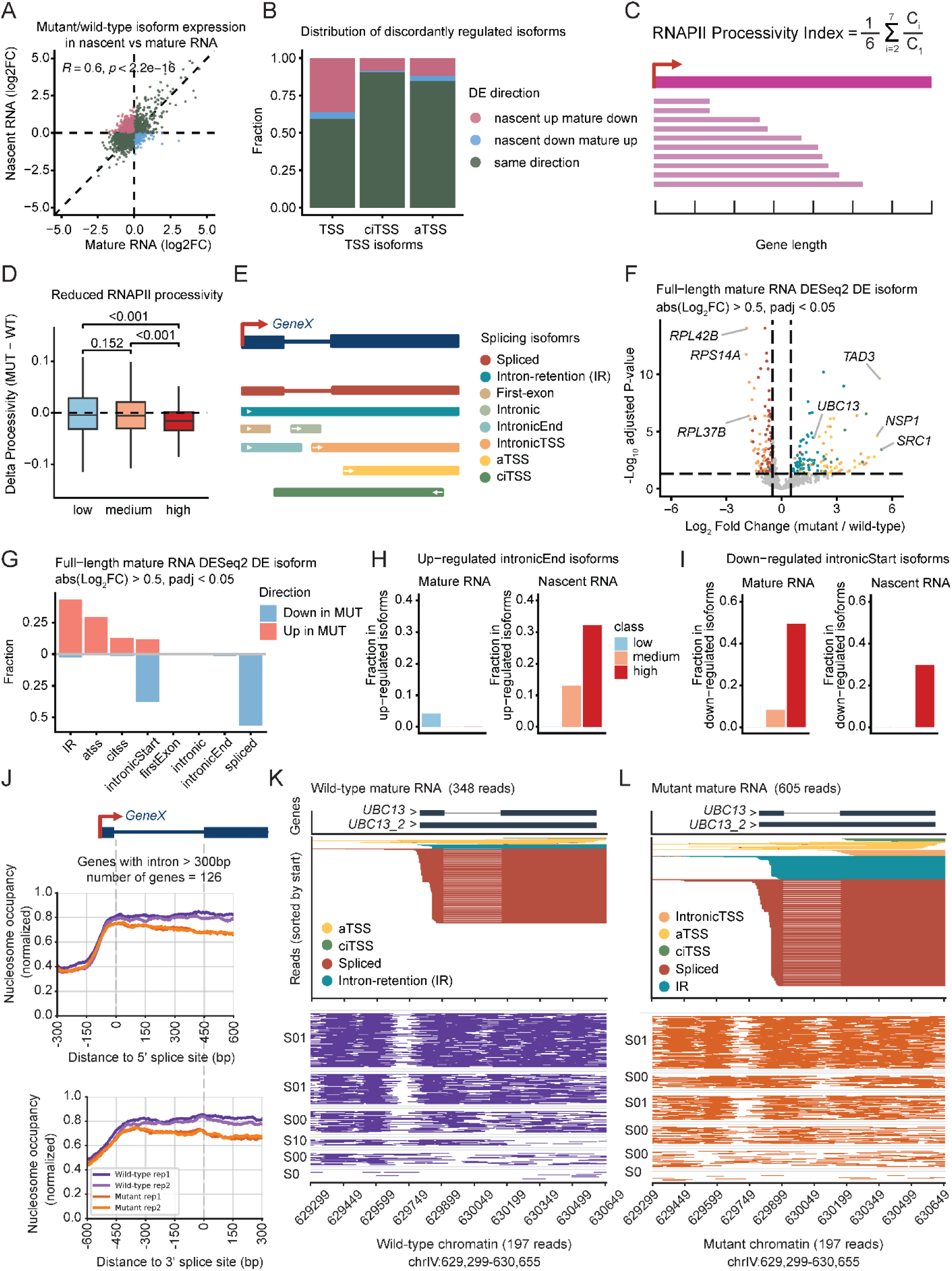
Loss of nucleosome spacing impairs RNAPII processivity and co-transcriptional splicing. (A) Scatter plot comparing log_2_ fold change (mutant vs wild-type) of TSS isoform expression between nascent RNA (y-axis) and full-length mature RNA (x-axis). Each point represents one isoform colored by the direction of discordance: nascent up and mature down (pink), nascent down and mature up (blue), or same direction in both libraries (teal). Pearson correlation coefficient and p-value are shown. (B) Stacked bar plot showing the fraction of TSS isoforms with concordant or discordant regulation between nascent and full-length mature RNA, stratified by TSS class. (C) Schematic illustrating the RNAPII processivity index calculation. Each gene body is divided into seven equal-length bins; the index is defined as the mean normalized read coverage across bins 2-7 relative to bin 1, capturing the degree to which nascent RNA coverage is maintained across the full gene body. Created in BioRender. Bai, G. (2026) https://BioRender.com/edm3lbt. (D) Boxplots of the change in RNAPII processivity index (delta processivity = mutant - wild-type) stratified by baseline expression class. Wilcoxon test p-values are shown for pairwise comparisons. Highly expressed genes show the greatest reduction in processivity in the *isw1*Δ *chd1*Δ mutant, indicating that remodeler loss disproportionately impairs elongation at the most actively transcribed loci. (E) Schematic illustrating the eight splicing isoform categories used to classify transcripts in intron-containing genes, based on the position of the TSS and TTS relative to the first exon and first intron: spliced, intron retention (IR), first-exon, intronic, intronicEnd, intronicTSS, aTSS, and ciTSS. Created in BioRender. Bai, G. (2026) https://BioRender.com/wnuaf5f. (F) Volcano plot of differentially expressed splicing isoforms between *isw1*Δ *chd1*Δ mutant and wild-type cells from full-length mature RNA sequencing, performed using DESeq2. Points above the significance threshold (|log_2_FC| > 0.5, padj < 0.05) are colored by splicing category. (G) Bar plot showing the fraction of differentially expressed splicing isoforms by category in full-length mature RNA, separated by direction of change (Up in MUT, salmon; Down in MUT, blue). (H) Bar plots showing the fraction of up-regulated intronicEnd isoforms among all up-regulated isoforms in full-length mature RNA (left) and nascent RNA (right), stratified by baseline expression class. (I) Bar plots showing the fraction of down-regulated intronicStart isoforms among all down-regulated isoforms in full-length mature RNA (left) and nascent RNA (right), stratified by baseline expression class. (J) Aggregate nucleosome occupancy profiles centered at the 5′ splice site (top) and 3′ splice site (bottom) for intron-containing genes with introns longer than 300 bp (n = 126 genes), shown for all four samples. (K-L) Genome browser track views of full-length mature RNA reads (top) and single-molecule chromatin fiber maps (bottom) at the *UBC13* locus (chrIV:629,299-630,655) in wild-type (K) and *isw1*Δ *chd1*Δ mutant (L) cells. RNA reads are colored by splicing isoform class.

The observed discrepancy in TSS isoform expression suggests an RNAPII processivity defect in the double mutant. To investigate this, we calculated an RNAPII processivity index by measuring the fraction of nascent RNA reads mapping to successive bins along each gene body (Figure 4C, Supplementary Table 7, Methods). Genes with high nascent transcription but low full-length mature RNA levels exhibited significantly reduced processivity in the mutant compared to other groups (Supplementary Figure 10C). Furthermore, when stratified by wild-type expression levels, genes with higher basal transcription showed a stronger correlation with processivity loss in the mutant (Figure 4D).

We next asked whether this processivity defect is driven by collisions between elongating RNAPII and polymerases newly initiating at cryptic intragenic TSSs. However, processivity loss was not associated with increased ciTSS expression (Supplementary Figure 10D, Supplementary Figure 10E, Supplementary Figure 10F), arguing against collision as the primary mechanism. Instead, processivity defects were most severe in highly expressed genes, which show the greatest disruption of gene-body nucleosome periodicity (Figure 1C) but the lowest levels of cryptic intragenic initiation (Figure 2D, Supplementary Figure 7B). Together, these observations suggest that the dominant mechanism of processivity impairment in the mutant is the direct disruption of regular nucleosome spacing, which in wild-type cells is thought to buffer torsional stress ahead of elongating RNAPII and facilitate polymerase passage through chromatin (Qian et al. 2026).

### Loss of nucleosome spacing increases intron retention and impairs co-transcriptional splicing

Co-transcriptional splicing is kinetically coupled to RNAPII elongation, and nucleosomes at splice sites are thought to facilitate this coupling (Carrillo Oesterreich et al. 2011; Lee et al. 2017; Wu et al. 2025). Given that RNAPII processivity is impaired in mutant cells, we next investigated whether co-transcriptional splicing was similarly affected. We analyzed intron-containing genes and classified nascent and mature RNA transcripts into eight categories based on splicing status and TSS position: antisense TSS (aTSS), cryptic intragenic TSS (ciTSS), firstExon, intronic, intronicEnd, intronicStart, intron retention (IR), and fully spliced isoforms (Figure 4E, Methods). Beyond the up-regulation of ciTSS and aTSS isoform expression, mutant cells showed a consistent decrease in spliced isoforms and a corresponding increase in IR isoforms in both nascent and full-length RNA (Figures 4F, 4G, Supplementary Figures 11A-D, Supplementary Table 8, 9).

Mutant cells exhibited an enrichment of intronicEnd nascent RNA transcripts, predominantly in highly expressed genes (Figure 4H, Supplementary Figure 11E, 11F). This accumulation implies an increase in partially processed nascent transcripts in the mutant. Furthermore, full-length mature RNA libraries revealed increased intron retention (IR) isoforms preferentially in medium- and high-expression genes (Figure 4I, Supplementary Figure 12A, 12B), suggesting that transcripts remained unspliced despite reaching the 3′ end. Collectively, these data demonstrate that the *isw1*Δ *chd1*Δ double mutant impairs splicing efficiency, with long-read sequencing of nascent and mature RNA revealing distinct populations of intron-retained transcripts (Supplementary Figure 12F).

This splicing defect is independent of antisense transcription (Supplementary Figure 12C-E) and is more pronounced in highly expressed genes (Supplementary Figure 12G), which showed the least cryptic transcription in the mutant. These results suggest that chromatin alterations are a more direct driver of the observed splicing phenotype. To investigate if chromatin alterations contribute to these splicing defects, we examined nucleosome occupancy at splice sites. Aggregated profiles centered on 5′ and 3′ splice sites showed reduced nucleosome occupancy in the mutant, with a more pronounced depletion at the 3′ splice site (Figure 4J). Accordingly, genes with significant splicing defects often display altered chromatin states, particularly enriched with co-accessible (S00) chromatin states (Supplementary Figure 12H). At single loci, in wild-type cells, splice sites are associated with well-positioned nucleosomes, whereas in the mutant, disrupted spacing increases splice site accessibility and correlates with elevated intron retention (Figure 4K, 4L, Supplementary Figure 11E, 11F). Together, these findings indicate that regular nucleosome spacing and splice-site nucleosome positioning are crucial for efficient co-transcriptional splicing, and that loss of gene-body nucleosome organization likely impairs spliceosome recruitment and catalysis.

## DISCUSSION

In this study, we integrated long-read chromatin profiling, nascent RNA sequencing, and full-length mature RNA sequencing to address a question that has been technically intractable with short-read approaches: are chromatin states associated with transcriptome output at the isoform level? Our findings reveal several interconnected results (Figure 5A). First, loss of *ISW1* and *CHD1* drives nucleosome redistribution from gene bodies toward promoter regions, reducing promoter accessibility while generating intragenic NDRs on the same chromatin fiber. Second, this coordinated chromatin reorganization coincides with a shift in isoform usage, decreased transcription from canonical TSSs and increased transcription from ciTSSs and aTSSs, establishing a direct link between single-molecule chromatin state and isoform output. Third, cryptic initiation is not random: ciTSSs are enriched for sequence motifs characteristic of canonical TSSs, including the Initiator element, and are flanked by well-positioned +1 and +2 nucleosomes that mirror canonical promoter architecture, indicating that cryptic initiation arises from pre-existing promoter-competent sequences normally suppressed by remodeler-maintained nucleosome occupancy. Fourth, intragenic nucleosome loss in the mutant is concentrated at ciTSS sites rather than distributed uniformly across gene bodies. Together, these findings support a unifying conclusion: nucleosome redistribution following remodeler loss decreases expression differences not only between canonical and cryptic TSSs within individual genes, but also between canonical TSSs of different genes, compressing the dynamic range of the transcriptional landscape toward a more uniform baseline.

**Figure 5.**
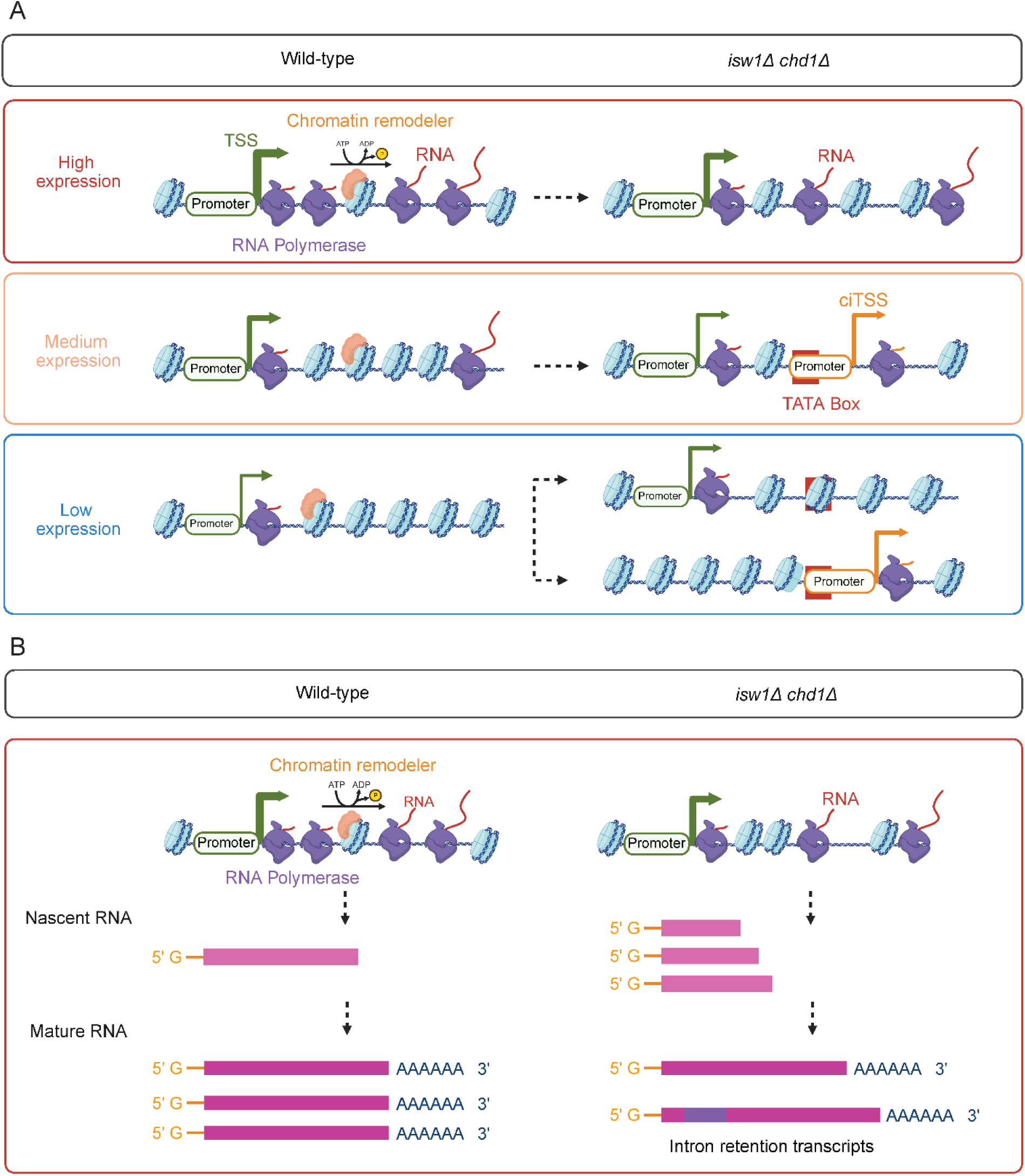
Model for *ISW1 and CHD1* remodeler-dependent maintenance of transcriptional fidelity at initiation and elongation. (A) Schematic model illustrating baseline transcription-dependent effects of *ISW1* and *CHD1* loss on transcription initiation. In wild-type cells (left), chromatin remodelers actively position nucleosomes to suppress promoter-competent sequences within gene bodies, suppressing cryptic transcription initiation. In *isw1*Δ *chd1*Δ mutant cells (right), loss of intragenic nucleosome exposes cryptic promoter-competent sequences in a transcription-dependent manner. In high-expression genes (red), disrupted nucleosome spacing does not lead to cryptic initiation, owing to the intrinsic depletion of TATA box elements within their gene bodies and/or competitive upstream TSS expression. In medium-expression genes (orange), exposed intragenic promoter elements activate ciTSS expression and suppress TSS expression, producing isoform switching from canonical to cryptic transcripts. In low-expression genes (blue), remodeler loss de-represses both TSS and cryptic TSSs, leading to broad transcriptional activation. Created in BioRender. Bai, G. (2026) https://BioRender.com/o9lcem7. (B) Schematic model illustrating the role of regular nucleosome spacing in supporting RNAPII processivity and co-transcriptional splicing. In wild-type cells (left), chromatin remodelers maintain evenly spaced nucleosome arrays that facilitate RNAPII elongation through the gene body, enabling the production of full-length, 5′-capped and polyadenylated mature transcripts. In *isw1*Δ *chd1*Δ mutant cells (right), disruption of nucleosome spacing impairs RNAPII processivity, resulting in accumulation of shorter nascent RNA reads and an increased abundance of intron-retention transcripts in the full-length mature RNA pool. Created in BioRender. Bai, G. (2026) https://BioRender.com/o9lcem7.

A profound consequence of *ISW1* and *CHD1* remodeler loss in our data is the compression of transcriptome expression variance: transcriptionally silent regions become de-repressed, whereas highly active transcription is reduced, flattening the transcriptional landscape toward a more uniform baseline. This pattern is consistent with the idea that ATP-dependent remodelers amplify differences in transcriptional output by establishing chromatin states that are more distinct than would arise from DNA sequence preferences alone (Shelansky et al. 2024). In particular, de-repression appears to result from increased exposure of Initiator (Inr) motifs at both canonical and intragenic sites following the loss of remodeler-imposed nucleosome occupancy, leading to the formation of promoter-like chromatin architecture. These promoter-competent sequences are present throughout the genome but normally suppressed, suggesting that remodeler activity, not DNA sequence alone, determines the extent of initiation.

Interestingly, although highly expressed genes show the greatest nucleosome disruption in the mutant, they exhibit relatively low levels of cryptic initiation. Previous studies have shown that strong upstream promoters can suppress the usage of downstream alternative promoters (Lee et al. 2021). Second, our motif analysis reveals that gene bodies of highly transcribed genes are intrinsically depleted of TATA box elements, likely reflecting negative selection against promoter-competent sequences in regions subject to frequent nucleosome disassembly during active transcription. Similarly, TATA boxes were found to be enriched in the promoters of up-regulated cryptic TSSs in Spt6 mutant yeast cells (Doris et al. 2018). Collectively, these findings indicate that chromatin remodeler-mediated nucleosome organization and intrinsic sequence composition together function safeguards against cryptic transcription.

Beyond its role as a transcription barrier to suppress transcription, our data also suggest that regularly spaced nucleosome arrays actively support RNAPII elongation and co-transcriptional splicing (Figure 5B). In wild-type cells, regular nucleosome spacing facilitates RNAPII transit through gene bodies, promoting production of full-length mRNAs. In the mutant, disruption of this spacing impairs RNAPII processivity in a manner that scales with baseline expression level: highly expressed genes, which are most dependent on remodeler activity to maintain nucleosome organization, exhibit the strongest processivity defects. Since processivity loss is not correlated with ciTSS or aTSS expression, collision between elongating and newly initiating RNAPII complexes is less likely to be the primary mechanism. Instead, processivity loss is due to direct impairment of RNAPII bypass at irregularly spaced nucleosomes, which in wild-type cells buffer torsional stress generated during elongation to facilitate polymerase passage (Qian et al. 2026).

The consequences of nucleosome spacing disruption extend beyond elongation to co-transcriptional splicing. Loss of *ISW1* and *CHD1* increased intron retention in both nascent and full-length RNA. Intriguingly, although slower RNAPII elongation is generally thought to favor splicing by providing more time for spliceosome assembly (Dujardin et al. 2014; Eick and Geyer 2013), our data show that nucleosome loss at splice sites is associated with decreased splicing efficiency. This apparent paradox suggests that the relevant parameter may not be elongation speed, but RNAPII stalling and loss of nucleosome occupancy at splice sites. Consistent with recent reports that nucleosomes and histone modifications such as H3K4me3 at the 3′ splice site facilitate the transit of splicing factors from RNAPII to the nascent transcript (Wu et al. 2025), our findings suggest that remodeler-maintained nucleosome positioning in intron-containing genic regions is required for efficient co-transcriptional splicing.

A key advantage of long-read chromatin profiling is the ability to measure promoter and intragenic chromatin states simultaneously on the same DNA molecule. Our data reveal that remodeler loss does not simply disrupt gene-body nucleosome positioning in a uniform way. Rather, the relationship between promoter and intragenic accessibility is actively rewired in an expression-dependent manner. More importantly, long-read chromatin fibers distinguish co-accessible states (S00) from mixed promoter-accessible (S01) or intragenic-accessible (S10) states, a distinction often obscured by aggregate short-read accessibility signals, thereby providing a more granular view of chromatin structure alterations.

Finally, our study establishes long-read multi-omic profiling as a powerful framework for directly connecting chromatin organization to isoform-level gene regulation. The central finding that the distribution of single-molecule chromatin states predicts isoform composition, in an expression-dependent manner and modulated by chromatin remodeler activity, extends our understanding of what chromatin organization encodes beyond the simple association between promoter accessibility and gene expression. Notably, loss of chromatin remodelers frequently reshapes isoform expression without proportional changes in total gene expression, demonstrating that the transcriptome is regulated at a finer grain than what is captured by gene-level quantification. This has important implications for interpreting chromatin perturbations. In diseases characterized by aberrant chromatin regulation, including cancers and developmental disorders, substantial layers of regulatory disruption may be overlooked if isoform composition is not analyzed alongside gene expression levels. More broadly, our findings suggest that understanding transcriptional regulation requires moving beyond population-averaged, gene-level measurements toward single-molecule approaches that resolve coordinated chromatin and RNA states, which ultimately determine transcript output.

## METHODS

### Yeast strains and genotype construction

#### Wild-type (ys18): MATalpha his3-11,15 leu2-3,112 can1-100 ura3Δ5

Mutant (yM209.13): *MATalpha his3-11,15 leu2-3,112 can1-100 ura3Δ5 PHO5[GC] chd1Δ::KanMX4 isw1Δ::URA3*

### SacCer3 gene annotation

We used the Saccharomyces cerevisiae version 13 (ares v13) gene annotation, developed in the Manny Ares laboratory at the University of California, Santa Cruz ([CSL STYLE ERROR: reference with no printed form.]). This annotation encompasses 6,685 genes and 6,980 transcript isoforms, including 6,590 intronless and 302 that contain one or more introns (Supplementary Figure 1A). The median transcript length is 1,038 base pairs (Supplementary Figure 1B).

### Short-read total RNA extraction and sequencing

Total RNA was prepared from logarithmically growing wild-type and mutant yeast cells (∼1.2e+7) in YPD. Hot acid phenol was used to extract total RNA following the protocol outlined by Green and Sambrook (Green and Sambrook 2021). One fourth of the total RNA was treated with DNase I (Turbo DNase I, Thermo Fisher) and cleaned on a ZymoResearch RNA Clean and Concentrate column. RNA yields were measured on a Qubit fluorometer, and a fraction of the RNA was also run on an Agilent TapeStation to assess RIN quality. Strand-specific Illumina bulk RNA sequencing of three biological replicates each from wild-type and mutant total RNA was performed by GeneWiz (Azenta Life Sciences) on Illumina NovaSeq 2x150bp.

### Short-read total RNA preprocessing, mapping, and quantification

Reads were aligned to the Saccharomyces cerevisiae sacCer3 reference genome with aresv13 gene annotations using STAR v2.7.11b (Dobin et al. 2013). Specifically, the STAR genome index was built with --runThreadN 32 --runMode genomeGenerate --genomeFastaFiles sacCer3.fa --sjdbGTFfile sacCer3_ares_v13.gtf --genomeSAindexNbases 10 --sjdbOverhang 149. Read alignment was performed in two-pass mode to improve splice junction detection using the following parameters “--runThreadN 16 --twopassMode Basic --alignIntronMin 40 --alignIntronMax 2000 --alignMatesGapMax 4000 --alignSJDBoverhangMin 1 --sjdbOverhang 149 --outSJfilterIntronMaxVsReadN 2000 2000 2000 --outSAMtype BAM SortedByCoordinate --quantMode TranscriptomeSAM”. Aligned reads were then quantified with RSEM v1.3.0 (Li and Dewey 2011) using parameters “--no-bam-output --seed 12345 -p 16 --strandedness reverse --alignments --paired-end”. The sample-level output from RSEM is concatenated into gene- and isoform-level count matrices and transcripts per million (TPM) matrices using the R package tximport (Soneson et al. 2015).

### Filtering and classification of genes by baseline expression

To accurately stratify genes into low-, medium-, and high-expression groups based on short-read RNA-seq data, we applied a series of filtering steps to remove genes that could not be reliably quantified. Because mitochondrial DNA is not wrapped around nucleosomes, mitochondrial genes were excluded from downstream analyses. To ensure reliable mapping of reads, genes located in highly repetitive regions were also removed. We used Newmap v0.2 ([CSL STYLE ERROR: reference with no printed form.]) to identify uniquely mappable regions using a minimum read length of 800 bp, corresponding to the median long-read RNA length in our dataset. Only genes located within uniquely mappable regions were retained. Genes within 6 kb of chromosome ends were classified as subtelomeric and excluded. We further excluded genes with an effective gene length of zero as estimated by RSEM, representing transcripts too short to be reliably quantified by short-read sequencing. Tandem and overlapping genes were removed because short-read data cannot unambiguously distinguish their expression. Small nuclear RNAs (snRNAs) were also excluded, as some are not polyadenylated or transcribed by RNA polymerase II and may therefore not be regulated by chromatin remodelers.

After filtering, the final annotation comprised 6,016 genes and 6,292 isoforms for downstream analyses. For gene expression stratification, TPM values were averaged across three wild-type replicates for each gene. Genes with an average TPM of zero were classified as low-expression. The expressed genes were ranked by average TPM, and expression percentiles were used to define categories: genes in the bottom 20% were assigned to the low-expression group, genes in the top 20% to the high-expression group, and genes in the middle 60% to the middle-expression group. In total, we defined 1,410 low, 3,453 medium, and 1,153 high-expression genes. This stratification allowed for systematic comparisons of chromatin and transcriptional alterations across expression categories.

### Short-read total RNA differential gene expression analysis

Differential gene expression analysis was performed using DESeq2 in R (Love et al. 2014). Genes were considered significantly differentially expressed if they met the criteria of an adjusted p-value < 1 × 10⁻³ and an absolute log₂ fold change (log₂FC) > 0.5. Significant genes were classified as Up in mutant (log₂FC > 0.5, FDR < 1 × 10⁻³) or Down in mutant (log₂FC < −0.5, FDR < 1 × 10⁻³). The top 2 differentially expressed genes were selected based on ranked log₂ fold change.

### Purification of Protein A-Hia5 (pA-Hia5)

pA-Hia5 was purified following the protocol of Altemose et al. (Altemose et al. 2021) with some modifications. *E. coli* cells were lysed with a single pass through an FA-031 pressure cell loaded on a French press (SLM Aminco, USA). Following nickel affinity purification (BioRad, Profinity), the pA-Hia5 sample was desalted into Buffer A (50mM Tris, 8.0, 100mM NaCl, 1mM DTT) using a 26/10 desalting column (Cytiva 17508701) and loaded onto tandem HiTrap Q/SP HP (Cytiva, 29951325 and 17115201) columns. Since pA-Hia5 binds poorly to either exchanger, the flow through and column wash (where >85% of the protein was found) were collected and run through a second nickel affinity column, followed by desalting into Buffer A with a PD10 G25 Desalting column (Cytiva). The enzyme was concentrated on an Amicon Ultra15 (10K cutoff) centrifugal unit, and activity, as described by Stergachis et al. 2020 (Stergachis et al. 2020), was measured.

### Nuclei isolation, methylation, and HMW DNA extraction and long-read DNA sequencing

Nuclei were isolated from wild-type and mutant cells according to the protocol outlined by Brahma and Henikoff (Brahma and Henikoff 2022). Six aliquots of nuclei were prepared from 200mL of logarithmically growing yeast cells in YPD (∼3.2e+9 cells), flash frozen in liquid nitrogen, and stored at -80 °C.

Two aliquots of nuclei (∼8e+8) were rapidly thawed for methylation by pA-Hia5. Briefly, nuclei were spun at 4,000 xg for 2 min and permeabilized in Digitonin buffer (20mM Hepes-KOH, 7.5, 150mM NaCl, 1mg/mL BSA, 0.15mM Spermine, 0.5mM Spermidine, 1x Roche Protease Inhibitors, 0.02% Digitonin) for 10min on ice and briefly spun at 4,000 xg for 2 min at 4 °C. The permeabilized nuclei were each resuspended in 750 μL of 1x Hia5 Sucrose Activation Buffer (15mM Tris-HCl, 8.0, 15mM NaCl, 60mM KCl, 1mM EDTA, 0.5mM EGTA, 0.34M Sucrose, 1mg/mL BSA, 0.5mM Spermidine, 0.15mM Spermine). Approximately 1,000U of Protein A-Hia5 were added to each aliquot of nuclei, mixed by inversion, and incubated on ice for 5min. The MTase reaction was started by the addition of a 1/40^th^ volume of 32mM SAM and incubated at 37 °C for 20min. A second aliquot of SAM was added after a 10min incubation. The reaction tubes were plunged into an ice-water bath for 2min before spinning the nuclei out at 4,000 xg/2min at 4 °C, and resuspending the nuclei in 40 μL 1x PBS. HMW DNA was extracted using the NEB Monarch HMW DNA extraction kit, quantified using the Qubit broad range DNA kit, and analyzed on an Agilent TapeStation tape. 4-5ug of HMW DNA was used for making libraries with the ONT SQK-LSK110 kit.

### Long-read DNA modification calling and nucleosome calling

DNA molecules were base-called with Dorado’s dna_r10.4.1_e8.2_400bps_sup@v4.2.0 simplex model, and modified bases were detected using Dorado’s dna_r10.4.1_e8.2_400bps_sup@v4.2.0_6mA@v2 model. Secondary and supplementary reads were filtered with samtools v1.3.1 (Li et al. 2009). Single-molecule nucleosome occupancy was predicted with fibertools-rs v0.6.2 (Jha et al. 2024) using the default parameters with “ft add-nucleosomes” followed by “ft extract”. The predicted nucleosomes are stored in a bed12 format. Reads aligned to the rDNA locus and mitochondrial DNA were filtered out from the downstream analysis.

### Long-range chromatin state clustering, annotation, and quantification

A long-read chromatin analysis workflow was built in EpiFLAIR. For each gene in the aresv13 annotation, chromatin reads spanning the genic region from upstream 500 bp of the TSS to downstream 50 bp of the TTS were extracted. Nucleosome occupancy matrices were constructed for each gene, where values of 1 indicate nucleosome-occupied positions, 0 indicate accessible positions, and −1 indicates missing data. Reads with missing values were filtered based on a user-defined fraction na_frac. Here, we used the default value na_frac=0 to only analyze chromatin that covers the entire region. Hypermethylated reads were removed if more than 60% of positions were marked as modified (mod_frac = 0.6). Genes with fewer than 10 remaining chromatin fibers were excluded from downstream analyses.

To cluster chromatin fibers for each gene, pairwise distances between reads were computed based on their nucleosome occupancy profiles. Specifically, the absolute difference between all read pairs was calculated and convolved along the genomic axis with a sliding window (default 150 bp) to emphasize regional run-length differences in nucleosome organization. The convolved differences were averaged across positions to generate an n × n distance matrix for n chromatin fibers. Hierarchical clustering was performed using agglomerative clustering with Ward linkage and Euclidean distance, and reads were partitioned into a fixed number of clusters (default n = 6).

For each cluster, a consensus nucleosome occupancy profile was computed by rounding the mean occupancy value at each position across all reads in the cluster. Continuous accessible segments (consensus value = 0) of at least 50 bp were defined as accessible regions. Using gene annotations, accessible regions were intersected with promoter, exon, and intron intervals to determine chromatin accessibility states for each region. Promoters were classified as accessible if an accessible region overlapped the promoter by at least 50 bp (or the promoter length if shorter). Genic regions were classified as accessible if at least one exon contained an accessible region meeting the same criterion.

Each cluster was annotated into one of five chromatin states based on the co-accessibility of promoters and genic regions: promoter-accessible (S01) when the promoter is open but intragenic regions are closed; intragenic-accessible (S10) when the promoter is closed but at least one exon is accessible; co-accessible (S00) when both the promoter and intragenic regions are open; fully nucleosomal (S1) when both promoter and intragenic regions are closed; or fully accessible (S0) defined as clusters in which >90% of the genic region was accessible. Clusters sharing the same annotation within a gene were collapsed, and the number of chromatin fibers from each sample assigned to each chromatin state was counted to generate per-gene, per–chromatin-state abundance matrices. These abundance matrices were then normalized by the total number of chromatin fibers per gene to obtain the fraction of each chromatin state per gene. Quantification was performed separately for each sample and merged at the end.

### Differential enrichment analysis of chromatin states

To identify chromatin states differentially enriched between mutant and wild-type conditions, we performed a contingency table–based enrichment analysis at the level of individual nucleosome or chromatin-state units (hereafter referred to as “events”). For each gene, counts corresponding to each event were extracted separately for two biological replicates per condition (mutant_rep1, mutant_rep2; wild-type_rep1, wild-type_rep2). For each event, 2 × 2 contingency tables were constructed comparing the number of reads (or signal counts) assigned to the focal event versus all other events within the same gene, between one mutant replicate and one wild-type replicate. A pseudocount of 1 was added to all cells in the table to avoid zero counts.

Statistical significance was assessed using the chi-square test of independence (χ² test), and an odds ratio was calculated from the contingency table. Log₂-transformed odds ratios (log₂OR) were computed to quantify effect size, where positive values indicate enrichment in mutant and negative values indicate enrichment in wild-type. Because two biological replicates were available per condition, all pairwise replicate comparisons between mutant and wild-type (four total comparisons) were performed for each event. P-values from these comparisons were combined using Fisher’s method to generate a single combined p-value per event. The mean log₂OR across replicate comparisons was used as the summary effect size. The number of nominally significant pairwise comparisons (p < 0.05) was also recorded to assess consistency across replicates. To evaluate within-condition reproducibility, additional 2 × 2 contingency tables were constructed comparing the two wild-type replicates and the two mutant replicates separately. Fisher’s exact test was used for these within-condition comparisons, and corresponding p-values were calculated to detect replicate-specific variability. Multiple hypothesis testing correction was performed using the Benjamini-Hochberg procedure to control the false discovery rate (FDR). Adjusted p-values were calculated separately for (i) combined between-condition p-values, (ii) wild-type replicate comparisons, and (iii) mutant replicate comparisons.

To quantify the magnitude of differential usage, mean signal values were calculated across replicates for each condition, and the difference between mutant and wild-type (Δ = mean_mutant − mean_wild-type) was computed for each event. The final output table with the event name, gene name, mean log₂OR, Δ usage, p-value, and adjusted p-value was used for downstream analysis. This framework enables robust detection of chromatin-state enrichment differences between conditions while accounting for biological replication and controlling for multiple testing.

### Identification and annotation of +1 nucleosomes and NDRs at the single-molecule level

The position of the +1 nucleosome was determined for individual chromatin fibers using single-molecule nucleosome occupancy profiles. For each read, run-length encoding was applied to identify continuous nucleosome-occupied segments (value = 1) and accessible segments (value = 0).

We implemented a hierarchical strategy to identify the +1 nucleosome. First, accessible regions longer than 75 bp were defined as candidate nucleosome-depleted regions (NDRs). Reads lacking any accessible region were classified as fully nucleosomal, whereas reads with a single accessible region spanning>90% of the gene body were classified as fully accessible. For the remaining reads, promoter accessibility derived from cluster-level consensus profiles was used to guide +1 nucleosome detection. If a promoter-accessible region was present, the NDR corresponding to the promoter was identified, and the +1 nucleosome was defined as the first nucleosome immediately downstream of the promoter NDR. If no promoter-accessible region was detected, the +1 nucleosome was instead defined as the first nucleosome downstream of the annotated TSS.

To annotate the genomic context of +1 nucleosomes, their positions were compared with gene structural features. For intronless genes, +1 nucleosomes were annotated as near_tss if located within 50 bp of the TSS, or downstream otherwise. For intron-containing genes, +1 nucleosomes were classified as near_tss, donor_ss (near the 5′ splice site), first_intron, acceptor_ss (near the 3′ splice site), or downstream. Reads containing internal NDRs without a detectable promoter-associated NDR were assigned the annotation intragenic-accessible.

For each +1 nucleosome call, upstream NDR was calculated as the distance between the upstream boundary of the +1 nucleosome and the downstream boundary of the preceding nucleosome. For each gene in each sample, +1 nucleosome positions, +1 nucleosome annotations, NDR positions, and NDR lengths were reported at the single-molecule level.

### Annotation of +1 nucleosomes and NDRs at the gene level

Single-molecule level +1 nucleosome and NDR features were further summarized for each gene. Reads annotated as fully accessible, fully nucleosomal, or intragenic-accessible were excluded, and only reads with identifiable +1 nucleosome positions were retained. Genes supported by fewer than 5 valid reads were discarded, as there is no reliable support for +1 nucleosome positions. For each gene, the median +1 nucleosome position was calculated and written to a BED file, while single-molecule +1 nucleosome counts were aggregated into a BEDGraph track. To quantify +1 nucleosome positional variability, the interquartile range (IQR) and Shannon entropy were computed for +1 positions. NDR lengths corresponding to the same valid reads were extracted, and the median NDR length per gene was calculated.

### Calculation of aggregated nucleosome occupancy relative to the +1 nucleosome dyad

For aggregated nucleosome occupancy analysis, we used the gene-level +1 nucleosome identified from EpiFLAIR. For each gene, we calculated the average nucleosome occupancy for positions up to 1000 bp upstream and downstream relative to each +1 nucleosome dyad. Finally, the scores from an individual gene locus were averaged to generate genome-wide aggregated nucleosome occupancy scores at +1 nucleosome dyads. The aggregated nucleosome occupancy scores from each sample were normalized by the peak of +1 nucleosomes.

### Long-read nascent RNA sequencing

Nascent RNA sequencing was performed as described by Robik (Robik et al. 2025) in 2 biological replicates. cDNA derived from nascent RNA was sequenced on an R10 MinION using the SQK-NBD114.24 Oxford Nanopore cDNA sequencing kit. Reads were basecalled with Dorado v0.7.3 using the dna_r10.4.1_e8.2_400bps_sup@v4.2.0 model. Adapter sequences were identified and trimmed using Pychopper (edlib mode), and only reads containing adapters at both ends were retained.

### Long-read nascent RNA sequencing quality control

Because nascent RNAs were not enriched using a 5′-capped library preparation, we applied the following quality control procedure to identify and retain reads originating from capped transcripts. Trimmed reads were aligned to the SacCer3 reference genome using minimap2 v2.24 (Li 2018) with parameters “-ax splice:hq -t 16 --secondary=no -G 3k”.

Inspection of the aligned reads revealed two characteristic features. First, a subset of reads contained poly(A) tails, indicating carry-over of polyadenylated mature RNA. Second, and more importantly, the majority of reads initiating from annotated TSS carried a non-genome-encoded 5′ guanosine (G), consistent with the presence of an mRNA 5′ cap, a pattern confirmed by comparison with matched public CAGE-seq data. Because the 5′ cap marks bona fide transcription start sites, reads bearing this feature were considered to represent nascent transcripts with known sites of transcription initiation.

To exploit this feature for filtering, we retained the terminal G from the 5′ sequencing adapter during trimming. When combined with the native 5′ cap guanosine, authentic capped reads produce exactly two non-genome-encoded 5′ G bases that appear as soft-clipped bases in genomic alignments. Reads containing poly(A) tails were therefore discarded, and only reads with exactly two soft-clipped 5′ Gs were retained as high-confidence nascent transcripts.

In intron-containing genes with short first exons, we observed that minimap2 occasionally failed to produce correct spliced alignments, instead generating spuriously long 5′ soft-clipped regions that caused genuine capped reads to fail the filtering criteria above. To recover these reads, we performed a second-pass alignment of the filtered read set against transcript sequences of intron-containing genes. Reads containing exactly two non-genome-encoded 5′ Gs in this alignment were retained and merged with the first-pass dataset to produce the final set of high-confidence nascent RNA reads.

### Long-read full-length mature RNA sequencing

cDNA libraries from three biological replicates of mature wild-type and mutant total RNA were generated using the TeloPrime Full-length cDNA Amplification kit v.2 (Lexogen). The six cDNA libraries were amplified with 14 PCR cycles, purified on AmPure XP beads, barcoded, and sequenced on an R10 MinION using the SQK-NBD114.24 Oxford Nanopore cDNA sequencing kit. Reads were basecalled with Dorado v0.7.3 using the dna_r10.4.1_e8.2_400bps_sup@v4,2,0 model.

### Construction of an extended yeast transcriptome

For each gene in the yeast gene annotation BED file, we calculated its distance (D) to the upstream neighboring gene. For non-overlapping genes, the transcription start site (TSS) was extended upstream by min(D/2, 150 bp). For overlapping genes (where the TSS of gene A overlaps with another gene B), we calculated a modified distance (D′) defined as the distance between the TSS of gene A and the nearest boundary of gene B (either the TSS or the transcription termination site), depending on strand orientation. The TSS of overlapping genes was then extended by min(D′, 150 bp).

### Long-read RNA genomic and transcriptomic alignment

We noticed that some transcript start and end positions of full-length mature RNAs deviate from the aresv13 gene annotations. In addition, extensive isoform overlap in the yeast transcriptome complicates accurate assignment of short transcripts, particularly those from nascent RNA. To improve transcript assignment and quantification for both nascent and mature RNAs, we developed a pipeline using both genomic and transcriptomic alignments. QC-passed reads from each RNA sequencing library (nascent and full-length mature) were aligned to both the genome and the extended yeast transcriptome (Methods). Genomic alignment used minimap2 v2.24 with parameters “-ax splice:hq -t 16 --secondary=no -G 3k”. Transcriptomic alignment was first performed using minimap2 with parameters “-ax map-ont -t 20 --secondary=no --for-only” to restrict mapping to sense-oriented transcripts. Reads that failed to align in this step were subsequently re-aligned using the same parameters without --for-only, enabling recovery of antisense transcripts. Transcriptomic alignments from both steps were merged and used to assign each read to an annotated isoform. All alignment files were sorted and indexed using samtools v1.3.1 (Li et al. 2009).

### RNA isoform quantification in intronless genes

Isoform quantification was performed using transcriptome alignments. Transcriptomic alignments were used to assign each read to an annotated isoform for both full-length mature RNA and nascent RNA. Reads with mapping quality (MAPQ) <10 or with <30% of read length aligned to the transcriptome were excluded. Genomic alignments were then used to extract the precise 5′ (TSS) and 3′ (TTS) positions of each read at single-nucleotide resolution, taking the leftmost mapped position for plus-strand reads and the rightmost mapped position for minus-strand reads.

To characterize alterations in the transcription start site, we quantified three isoform types for each intronless gene based on the TSS position: TSS, cryptic intragenic TSS (ciTSS), and antisense TSS (aTSS) isoforms. For each gene, the 25th percentile of the gene length was used as a boundary to distinguish TSS (upstream of the 1st quantile) from ciTSS (downstream of the 1st quantile). Reads mapped to the reverse complement of the gene were assigned to the aTSS isoform. The quantification was performed on a per-sample basis and concatenated into a count table.

### Annotation of transcription start site in intronless genes

TSS positions from genomic alignments were recorded in strand-specific bedGraph format with TSS isoform annotation, yielding per-position read counts for each TSS, ciTSS, and aTSS isoform across all samples. BedGraph files were then collapsed to single representative TSS positions. For each isoform, all TSS positions weighted by read count were aggregated, and the modal TSS position was taken as the representative site. Isoforms supported by 3 or fewer reads were excluded to ensure reliability. The resulting per-isoform TSS positions were written to BED format with strand information retained. This classification was performed independently for each biological replicate and for both nascent and full-length mature RNA libraries, yielding per-replicate TSS, ciTSS, and aTSS BED files that were subsequently used as reference TSS coordinates.

### RNA isoform quantification in intron-containing genes

For intron-containing genes, isoform expression was quantified based on the splicing pattern of the first intron to represent variation in splicing and transcription initiation sites. Transcripts were classified as firstExon when the TTS was located within the first exon; intronicEnd when the TSS resided in the first exon and the TTS in the first intron; intronic when both TSS and TTS were located within the first intron; intronicStart when the TSS was within the first intron and the TTS within the second exon; intron-retention when the TSS was in the first exon and the TTS in the second exon with the first intron unspliced; and spliced when the first intron was fully removed. The quantification was performed on a per-sample basis and concatenated into a count table.

### Quantification of RNAPII Processivity Index

To quantify RNAPII processivity, each gene body was divided into seven equal-length bins, and nascent RNA read coverage was calculated per bin per gene. To minimize the contribution of reads originating from ciTSSs, which would artificially inflate coverage in internal bins and confound processivity estimates, reads were filtered to retain only those whose 5′ cap position fell within ±10 bp of the TSSs. Genes with fewer than 10 reads mapping to the first bin after filtering were further excluded to ensure reliable coverage estimates. For each retained gene, per-bin coverage was normalized to the coverage of the first bin, which carried the maximum coverage across all bins. The RNAPII processivity index was then defined as the mean of normalized coverage across bins 2 through 7, such that genes with uniform coverage across the gene body approach a value of 1 and genes with progressive 3′ coverage loss approach 0.

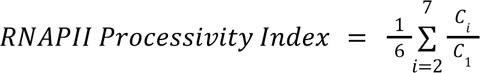

Processivity indices were calculated separately for each biological replicate and averaged within wild-type and mutant genotypes for downstream comparisons.

## CODE AVAILABILITY

Code for the execution of EpiFLAIR (https://github.com/baigal628/epiflair) and for reproducing analyses and figures in this manuscript (https://github.com/baigal628/saccer_lr_multiome_manuscript) is publicly available on GitHub.

## DATA AVAILABILITY

All raw and processed sequencing data generated in this study have been submitted to the NCBI BioProject database (https://www.ncbi.nlm.nih.gov/bioproject/) under accession number PRJNA1531117.

## ACKNOWLEDGMENTS

We thank Kousik Sundararajan (Department of Biochemistry, Stanford University) for advice on pA-Hia5 purification. We thank Manny Ares (Department of Molecular, Cell, and Developmental Biology, University of California, Santa Cruz) for providing the *Saccharomyces cerevisiae* gene annotation and for valuable scientific guidance. We thank Karla Neugebauer (Department of Molecular Biophysics and Biochemistry, Yale University) for sharing the nascent RNA sequencing protocol and for insightful discussions. We thank Tracy Jonsson (University of California, Los Angeles) for helpful scientific guidance. We are grateful to Carrie Partch (Department of Biochemistry, University of California, Santa Cruz) and Harry Noller (Department of Molecular, Cell, and Developmental Biology, University of California, Santa Cruz) for generously providing access to their lab equipment.

## AUTHOR CONTRIBUTIONS

Conceptualization and design were by A.N.B. and H.B. Experimental design and execution were by N.D. and B.M. Data analyses were by G.B., C.F., and H.H. Computational tool implementation was by G.B. Manuscript writing was by G.B. and N.D. with input from all authors.

## SUPPLEMENTARY TABLE LEGENDS

**Supplementary Table 1. Long-read chromatin sequencing quality control metrics.**

Per-sample sequencing statistics for long-read chromatin profiling. Columns report the total number of primary reads, the number of primary mapped reads, and the mean, median, and standard deviation of read length and read quality score.

**Supplementary Table 2. Differential chromatin state usage between wild-type and *isw1*Δ *chd1*Δ double mutant.**

Results of chromatin state differential enrichment analysis performed using the chi-square test across all genes and chromatin states. Each row corresponds to one chromatin state within one gene. Columns from left to right report chromatin state ID, gene ID, log_2_ odds ratio, combined p-value across replicate comparisons, number of nominally significant pairwise comparisons, within-condition p-values for wild-type and mutant replicates, Benjamini-Hochberg adjusted combined p-values, average chromatin state usage in mutant and wild-type, difference in chromatin state usage mutant - wild-type, chromatin state, differential enrichment label, and baseline expression class.

**Supplementary Table 3. Long-read nascent RNA sequencing quality control metrics.**

Per-sample sequencing statistics for long-read nascent RNA libraries. Columns report the total number of reads before filtering, reads removed due to absence of a 5′ cap signature, reads removed due to presence of a poly(A) tail, the number of high-confidence nascent RNA reads retained after filtering, the number of primary mapped reads, and the mean, median, and standard deviation of read length and read quality score.

**Supplementary Table 4. Long-read full-length mature RNA sequencing quality control metrics.** Per-sample sequencing statistics for full-length mature RNA libraries generated using the TeloPrime kit. Columns report the total number of primary reads, the number of primary mapped reads, and the mean, median, and standard deviation of read length and read quality score.

**Supplementary Table 5. Differential TSS isoform expression in nascent RNA.**

Results of differential expression analysis of TSS isoforms from nascent RNA sequencing, performed using limma. Each row corresponds to one isoform in one gene. Columns report isoform name, mean expression in wild-type and mutant, log_2_FoldChange, Benjamini-Hochberg adjusted p-value, differential expression label, gene ID, isoform ID, TSS annotation, and baseline expression class.

**Supplementary Table 6. Differential TSS isoform expression in full-length mature RNA.**

Results of differential expression analysis of TSS isoforms from full-length mature RNA sequencing, performed using DESeq2. Each row corresponds to one isoform in one gene. Columns report isoform name, the DESeq2 base mean expression, log_2_FoldChange, standard error of the log_2_FoldChange, Wald test statistic, p-value, Benjamini-Hochberg adjusted p-value, shrinkage-estimated log_2_FoldChange, differential expression label, gene ID, isoform ID, TSS annotation, and baseline expression class.

**Supplementary Table 7. RNAPII processivity index per gene.**

Per-gene RNAPII processivity indices calculated from nascent RNA sequencing for wild-type and *isw1*Δ *chd1*Δ mutant cells. Each row corresponds to one gene. Columns report sample processivity indices, averaged processivity indices per genotype, and the difference between mutant and wild-type. The processivity index is defined as the averaged nascent RNA read coverage across gene body bins 2-7, normalized to bin 1 coverage, such that values approaching 1 indicate uniform coverage and values approaching 0 indicate progressive 3′ coverage loss.

**Supplementary Table 8. Differential RNA splicing isoform expression in nascent RNA.**

Results of differential expression analysis of splicing isoforms from nascent RNA sequencing, performed using limma. Each row corresponds to one splicing isoform in one intron-containing gene. Columns report isoform name, mean expression in wild-type and mutant, log_2_FoldChange, Benjamini-Hochberg adjusted p-value, differential expression label, gene ID, isoform ID, splicing annotation, and baseline expression class.

**Supplementary Table 9. Differential RNA splicing isoform expression in full-length mature RNA.**

Results of differential expression analysis of splicing isoforms from full-length mature RNA sequencing, performed using DESeq2. Each row corresponds to one splicing isoform in one intron-containing gene. Columns report isoform name, DESeq2 base mean expression, log_2_FoldChange, standard error, Wald test statistic, p-value, Benjamini-Hochberg adjusted p-value, shrinkage-estimated log_2_FoldChange, differential expression label, gene ID, isoform ID, splicing annotation, and baseline expression class.

