## Supplementary Figures for "Multi-omic long-read sequencing reveals nucleosome control of transcriptional fidelity at the isoform level"

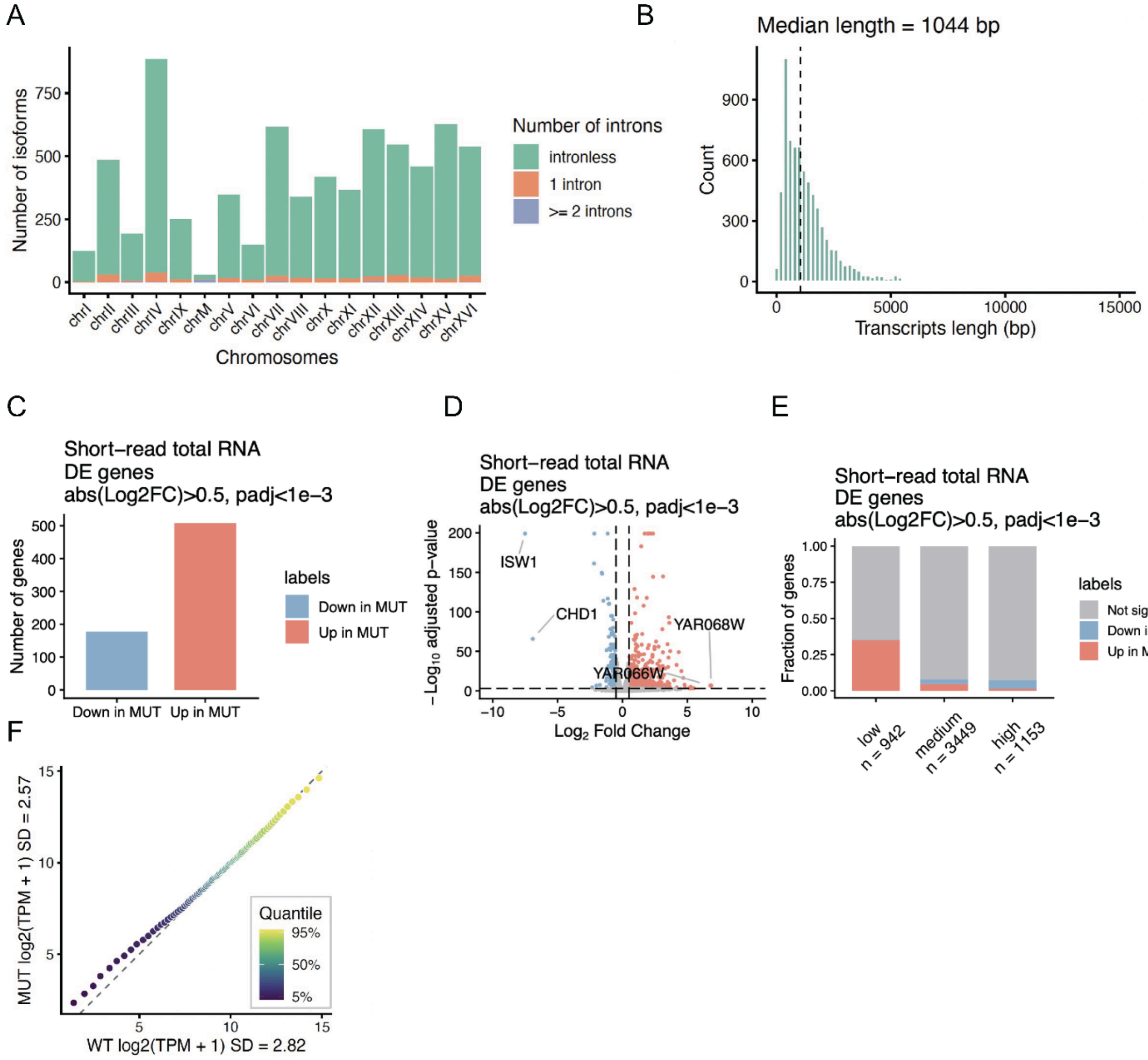


**Supplementary Figure 1. Gene annotation and short-read RNA-seq differential expression analysis in wild-type and *isw1*Δ *chd1*Δ mutant yeast.**

(A) Bar plot showing the number of isoforms per chromosome in the aresv13 S. cerevisiae gene annotation, colored by number of introns: intronless (teal), one intron (orange), and two or more introns (purple).

(B) Histogram of transcript lengths across all annotated isoforms. The dashed line indicates the median transcript length of 1,044 bp.

(C) Bar plot showing the number of differentially expressed genes between the *isw1*Δ *chd1*Δ mutant and wild-type cells identified by short-read total RNA-seq ($|\log_2 FC| > 0.5$, padj $< 1\times10^{-3}$).

(D) Volcano plot of differentially expressed genes from short-read total RNA-seq. Each point represents one gene. The top 2 up- and down-regulated genes *ISW1*, *CHD1*, *YAR066W*, and *YAR068W* are labeled.

(E) Stacked bar plot showing the fraction of genes in each expression class (low, medium, high) that are up-regulated, down-regulated, or not significantly changed in the mutant.

Low-expression genes are preferentially up-regulated, while high-expression genes are preferentially down-regulated.

(F) Scatter plot comparing gene-level expression (log2(TPM+1)) between wild-type (x-axis, SD (standard deviation) = 2.82) and *isw1*Δ *chd1*Δ mutant (y-axis, SD = 2.57), colored by expression quantile.

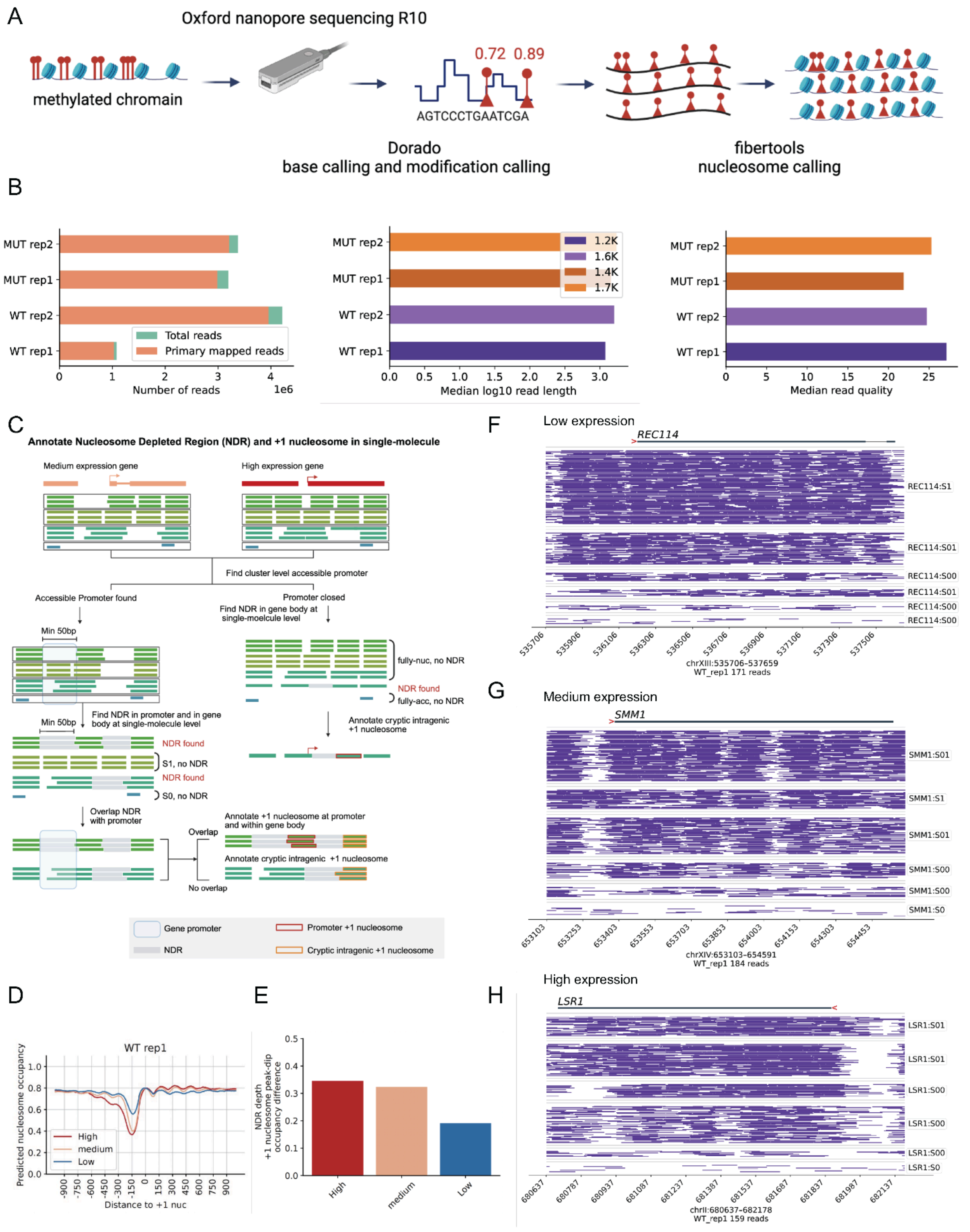


**Supplementary Figure 2. Long-read chromatin sequencing quality control and single-molecule nucleosome profiling in wild-type cells.**

(A) Schematic of the long-read chromatin accessibility sequencing workflow. pA-Hia5 methylated chromatin is sequenced on Oxford Nanopore R10 flow cells. The raw signal is basecalled, and modification called using Dorado, and nucleosome occupancy is predicted from individual reads using fibertools. Created in BioRender. Bai, G. (2026) https://BioRender.com/6lhglwt.
(B) Bar plots showing per-sample sequencing statistics for long-read chromatin libraries (2 wild-type replicates, 2 *isw1*Δ *chd1*Δ mutant replicates): total and primary mapped reads (left), median log10 read length (middle), and median read quality score (right).
(C) Schematic of the EpiFLAIR pipeline for annotating NDRs and +1 nucleosome positions at single-molecule resolution. Created in BioRender. Bai, G. (2026) https://BioRender.com/64bhnp2.
(D) Aggregate nucleosome occupancy profiles centered at the +1 nucleosome dyad for low-, medium-, and high-expression genes in wild-type rep1.
(E) Bar plot showing NDR depth measured by difference in peak-to-dip nucleosome occupancy stratified by baseline expression class. NDR depth scales progressively with transcriptional activity.
(F-H) Representative genome browser track views of single-molecule chromatin fibers for low: *REC114* (F), medium: *SMM1* (G), and high: *LSR1* (H) expression genes in wild-type rep1.

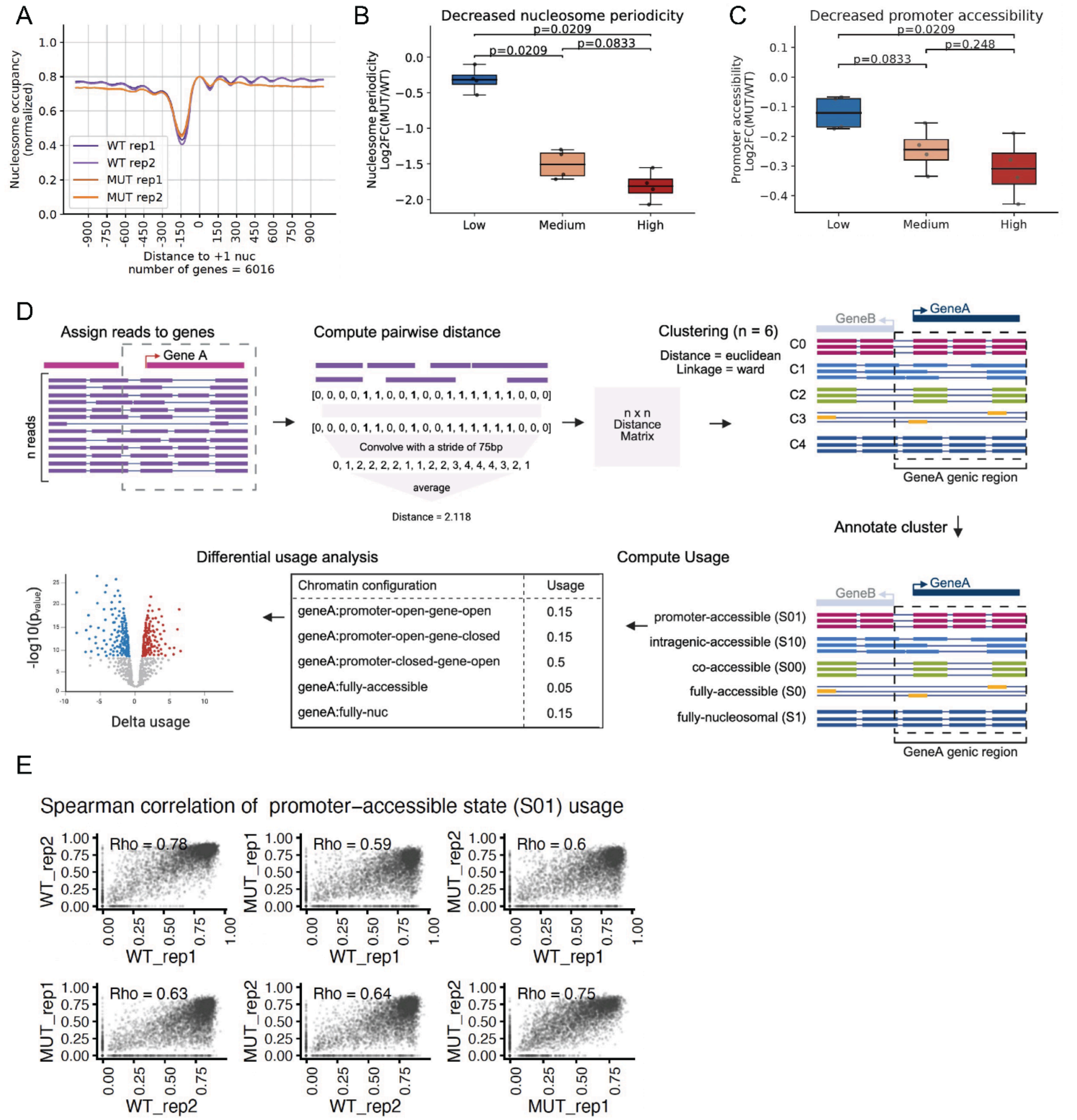


**Supplementary Figure 3. Analyses of single-molecule chromatin state in *isw1*Δ *chd1*Δ mutant cells.**

(A) Aggregate nucleosome occupancy profiles centered at the +1 nucleosome dyad for all 6,016 genes, comparing wild-type and *isw1*Δ *chd1*Δ mutant cells across 2 biological replicates.

(B) Boxplots showing log2 fold change (mutant vs wild-type) of nucleosome periodicity stratified by baseline expression class (low, medium, high), with Wilcoxon test p-values shown. Gene-body nucleosome periodicity is most severely disrupted in highly expressed genes.

(C) Boxplots showing log2 fold change (mutant vs wild-type) of promoter accessibility stratified by baseline expression class, with Wilcoxon test p-values shown. Promoter accessibility decreases most strongly in highly expressed genes.

(D) Schematic of the EpiFLAIR chromatin fiber clustering and chromatin state quantification workflow. Individual reads spanning each gene are clustered by pairwise nucleosome occupancy distance, annotated into discrete chromatin states, and summarized as per-gene chromatin state abundance matrices normalized by total fiber count. Created in BioRender. Bai, G. (2026) https://BioRender.com/kwiyip9.

(E) Scatter plots showing Spearman correlation of promoter-accessible state (S01) usage between wild-type and *isw1Δ chd1Δ* mutant replicates.

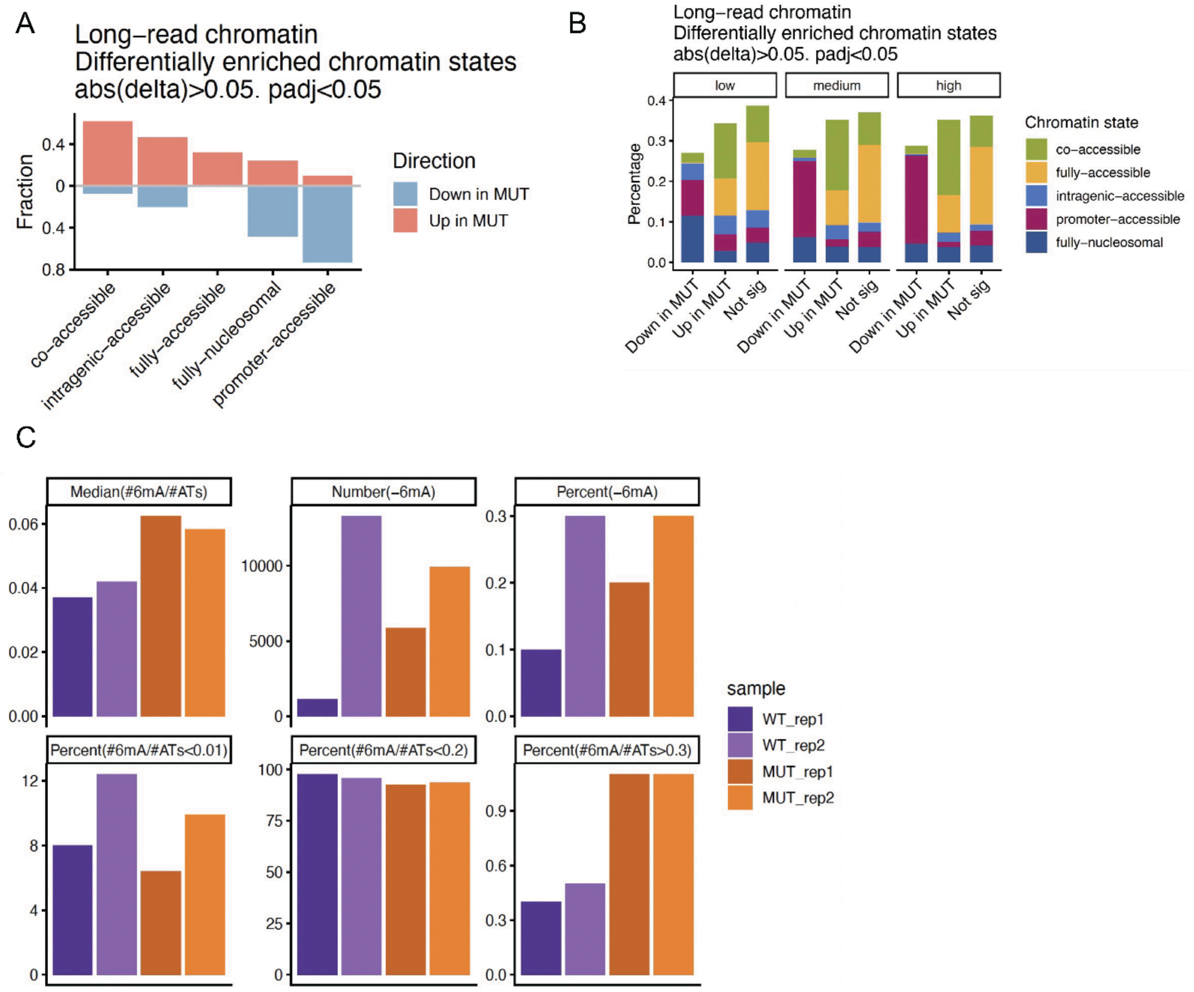


**Supplementary Figure 4. Global chromatin state changes and 6mA methylation quality control.**
(A) Bar plot showing the fraction of genes with each chromatin state significantly enriched (Up in mutant, salmon) or depleted (Down in mutant, blue) in the *isw1*Δ *chd1*Δ mutant relative to wild-type cells (|delta| > 0.05, padj < 0.05). The co-accessible (S00) and intragenic-accessible (S10) states are predominantly gained, while the promoter-accessible (S01) state is predominantly lost in the mutant.
(B) Stacked bar plots showing the proportion of each chromatin state among genes that are down-regulated, up-regulated, or not significantly changed in the mutant, stratified by baseline expression class (low, medium, high).
(C) Bar plots showing six per-sample metrics of 6mA methylation quality quantified by fiberseq-qc: median 6mA fraction per adenosine and thymine, number and percent of unmethylated adenosines, and fraction of reads with methylation below 0.01, below 0.2, or above 0.3.

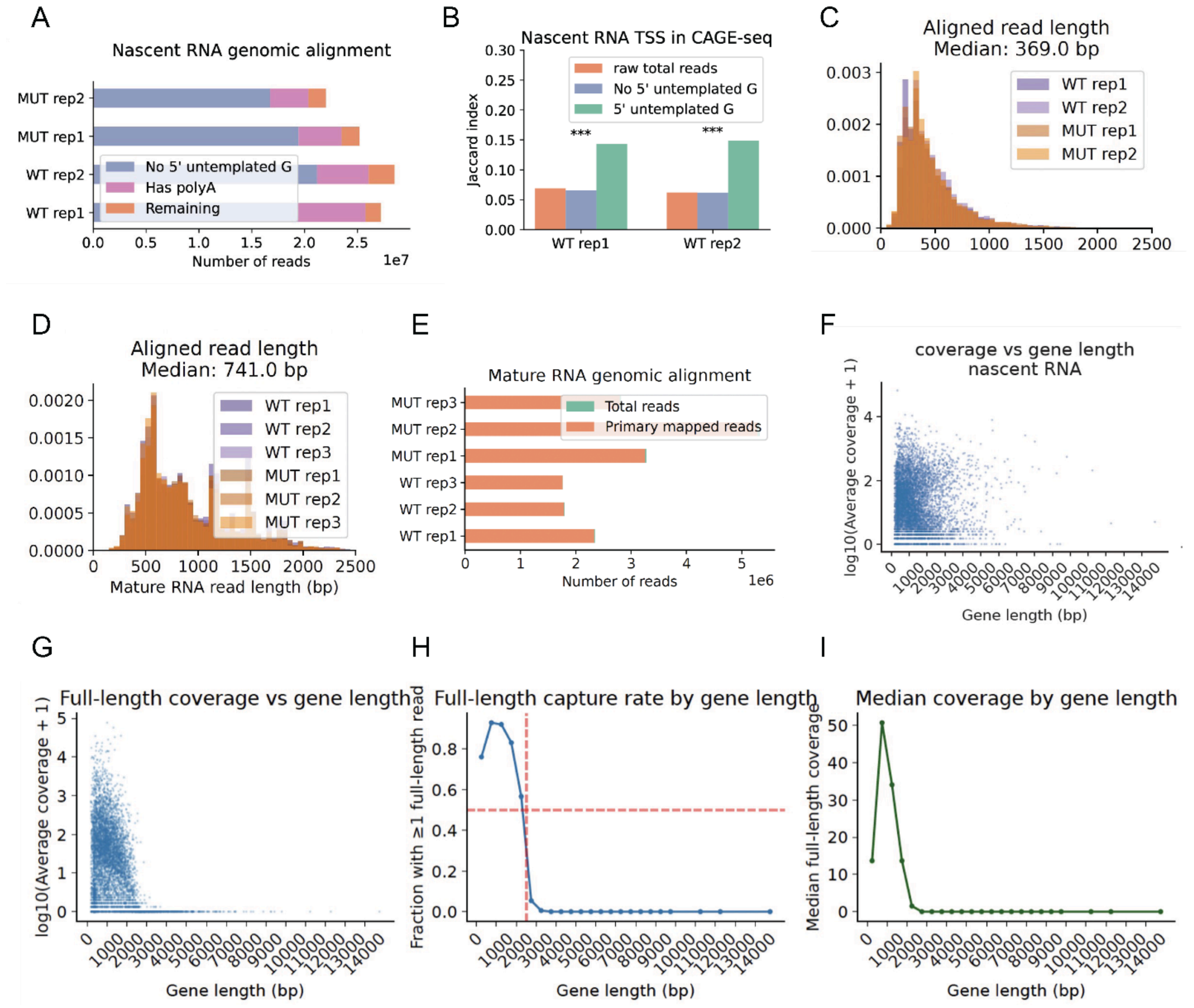


**Supplementary Figure 5. Quality control for long-read nascent and full-length mature RNA sequencing.**

(A) Bar plot showing the composition of nascent RNA reads per sample before and after filtering: reads lacking a 5' untemplated G (purple), reads carrying a poly(A) tail (pink), and reads retained as high-confidence capped nascent transcripts (orange).

(B) Bar plot showing Jaccard index between nascent RNA TSS positions and public CAGE-seq data for wild-type rep1 and rep2, comparing raw reads, reads without a 5' untemplated G, and reads with a 5' untemplated G. Reads carrying the 5' untemplated G signature show significantly higher overlap with CAGE-seq TSSs (*** $p < 0.001$), confirming that this signature identifies genuine capped transcripts.

(C) Histogram showing aligned read length of 5' capped nascent RNA reads across all four samples. Median aligned read length is 369 bp.

(D) Histogram of aligned full-length mature RNA read lengths across all six samples. Median aligned read length is 741 bp.

(E) Bar plot showing total and primary mapped reads per sample for the six full-length mature RNA libraries.
(F) Scatter plot of average nascent RNA coverage versus gene length, showing no systematic length bias in the nascent RNA library.
(G) Scatter plot of average full-length mature RNA coverage versus gene length, showing progressive coverage loss for longer genes due to reverse transcriptase processivity limitations.
(H) Line plot showing the fraction of genes with at least one full-length mature RNA read as a function of gene length. Genes shorter than ~2,000 bp have a capture rate above ~50%.
(I) Line plot showing median full-length mature RNA coverage as a function of gene length. Coverage drops sharply for genes longer than ~2,000 bp, confirming the read-length bias of the mature RNA library.

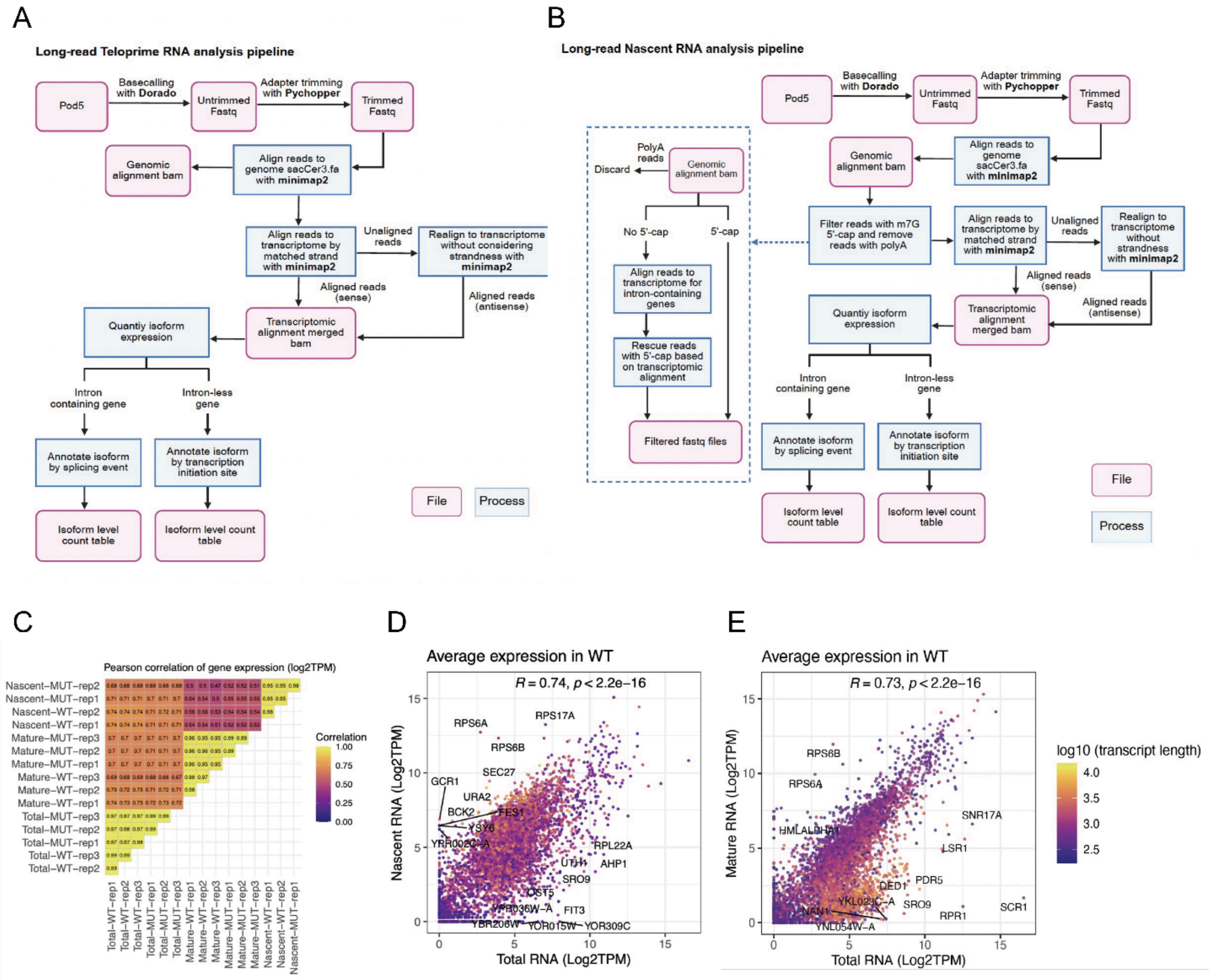


**Supplementary Figure 6. Long-read RNA isoform quantification pipelines and cross-library expression correlations.**

(A) Schematic of the long-read TeloPrime full-length mature RNA analysis pipeline, from basecalling with Dorado through adapter trimming with Pychopper, genomic and transcriptomic alignment with minimap2, and isoform-level quantification for intron-containing and intronless genes. Created in BioRender. Bai, G. (2026) https://BioRender.com/rr5dvfw.

(B) Schematic of the long-read nascent RNA analysis pipeline, including the 5' cap filtering step (dashed box) in which reads are filtered for the m7G cap signature and poly(A) tails are removed before alignment and isoform quantification. Created in BioRender. Bai, G. (2026) https://BioRender.com/jceja34.

(C) Heatmap of Pearson correlations of gene-level expression ($\log_2$TPM) between all pairwise combinations of nascent RNA, full-length mature RNA, and short-read total RNA samples. Expression estimates are more highly correlated within library type than between library types.

(D) Scatter plot comparing gene-level expression ($\log_2$TPM) between short-read total RNA (x-axis) and nascent long-read RNA (y-axis) in wild-type cells (Pearson R = 0.74, $p < 2.2\times10^{-16}$). Points are colored by log10 transcript length, and selected genes are labeled.

(E) Scatter plot comparing gene-level expression between short-read total RNA (x-axis) and full-length mature long-read RNA (y-axis) in wild-type cells (Pearson R = 0.73, $p < 2.2\times10^{-16}$). The lower correlation compared to nascent RNA is consistent with the read-length bias of the mature RNA library.

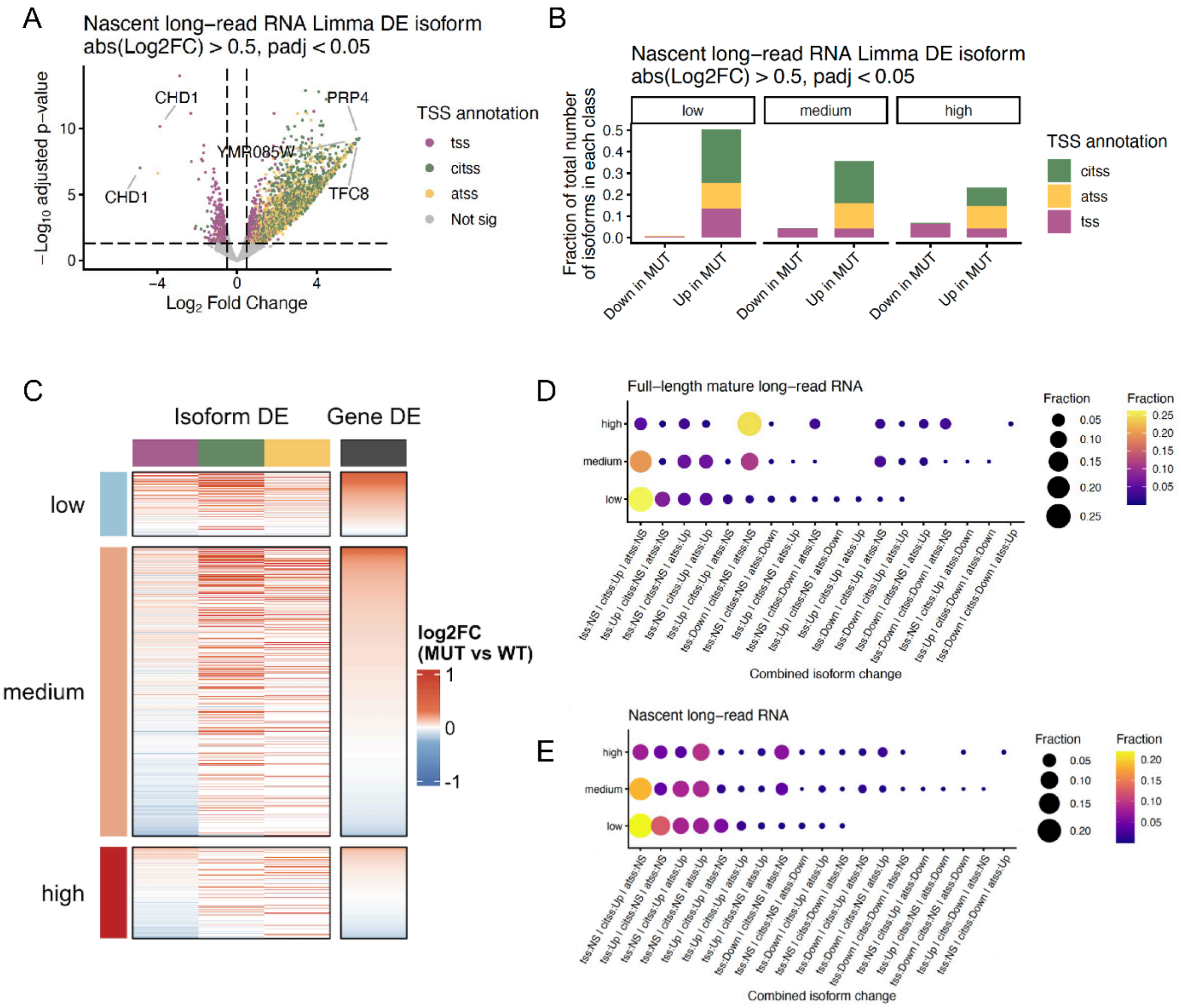


**Supplementary Figure 7. Differential TSS isoform usage in nascent RNA and co-occurring isoform change patterns.**

(A) Volcano plot of differentially expressed TSS isoforms between *isw1*Δ *chd1*Δ mutant and wild-type cells from nascent RNA. Each point represents one TSS isoform. Points above the significance threshold (|$\log_2$FC| > 0.5, padj < 0.05, limma) are colored by TSS annotation.

(B) Stacked bar plots showing the distribution of differentially expressed TSS isoforms from nascent RNA (|log2FC| > 0.5, padj < 0.05, limma), stratified by baseline expression class (low, medium, high) and direction of change, colored by TSS class (TSS, ciTSS, aTSS).

(C) Heatmap of isoform-level $\log_2$fold change (left, mutant vs wild-type) from nascent RNA and gene-level log2 fold change (right), stratified by baseline expression class. Each row represents one gene, illustrating that isoform-level changes frequently occur without proportional changes in total gene expression.

(D-E) Bubble plot showing the fraction of genes in each expression class with each combination of co-occurring TSS isoform changes (Down: down-regulated in the mutant, Up: up-regulated in the mutant, or NS: no significant change) in full-length mature RNA (D) and nascent RNA (E). Bubble size and color reflect the fraction of genes.

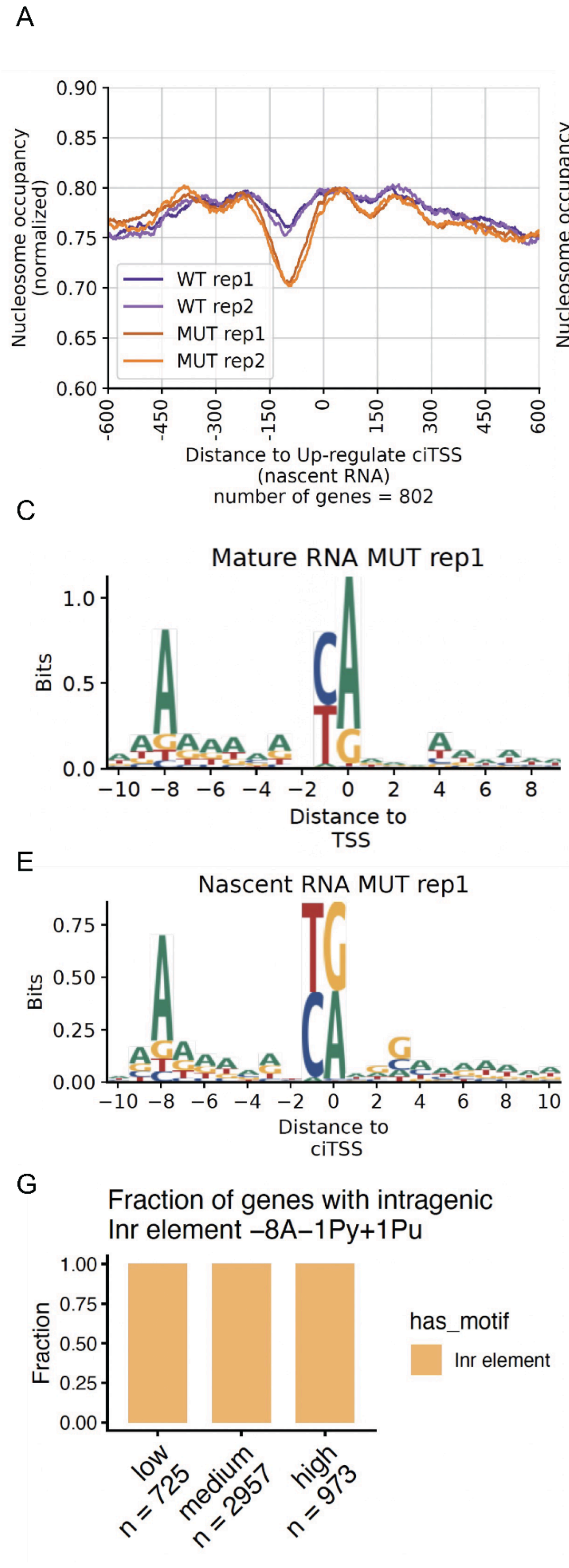

A
B
Nucleosome occupancy (normalized)
WT rep1
WT rep2
MUT rep1
MUT rep2
Distance to Up-regulate ciTSS (nascent RNA)
number of genes = 802
Distance to up-regulated aTSS (nascent RNA)
number of genes = 553
C
D
Mature RNA MUT rep1
Nascent RNA MUT rep1
Bits
Distance to TSS
E
F
Nascent RNA MUT rep1
Distance to ciTSS
Distance to aTSS
G
Fraction of genes with intragenic Inr element −8A−1Py+1Pu
Fraction
has_motif
Inr element
low n = 725
medium n = 2957
high n = 973

**Supplementary Figure 8. Chromatin architecture and sequence features at ciTSS and aTSS sites.**

(A) Aggregate nucleosome occupancy profile centered at up-regulated ciTSSs in the *isw1Δ chd1Δ* mutant from nascent RNA across all four samples (n = 802 genes). An NDR flanked by well-positioned +1 and +2 nucleosomes mirrors the chromatin architecture of canonical promoters.

(B) Aggregate nucleosome occupancy profile centered at up-regulated aTSSs in *isw1Δ chd1Δ* mutant cells from nascent RNA across all four samples (n = 553 genes). An NDR flanked by well-positioned +1 and +2 nucleosomes mirrors the chromatin architecture of canonical promoters.

(C-D) Sequence logos of the Initiator (Inr) motif centered at TSSs in full-length mature RNA (C) and nascent RNA (D) from *isw1Δ chd1Δ* mutant replicate1, showing the conserved PyPu dinucleotide at −1/+1 with an A at −8.

(E-F) Sequence logos of the Inr motif at ciTSSs (E) and aTSSs (F) from nascent RNA in the *isw1Δ chd1Δ* mutant rep1.

(G) Bar plot showing the fraction of genes in each expression class (low n=725, medium n=2957, high n=973) containing an intragenic Inr element (−8A−1Py+1Pu).

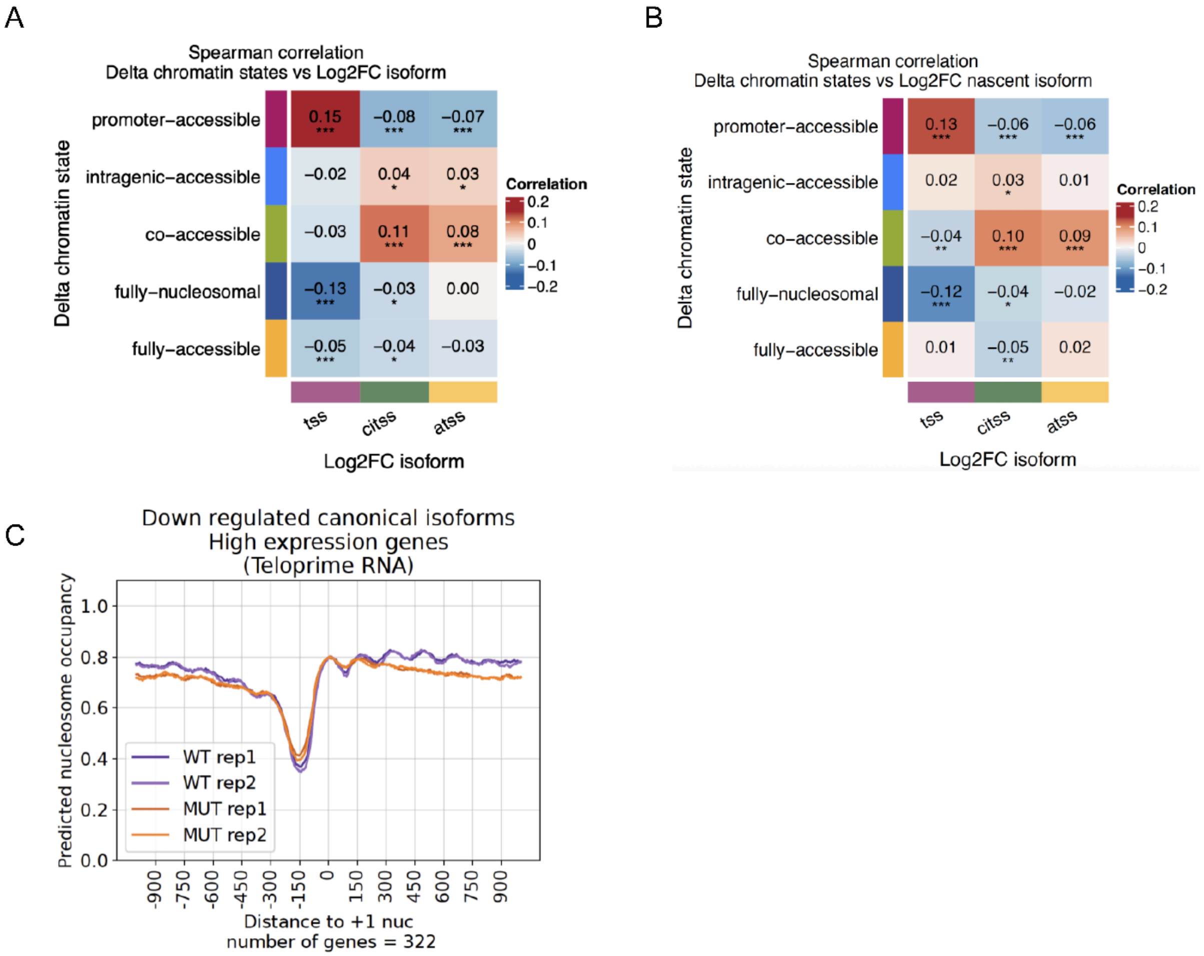


**Supplementary Figure 9. Correlation between chromatin state abundance change vs isoform expression change.**

(A-B) Heatmap of Spearman correlations between changes in chromatin states (Δ fraction = mutant - wild-type) and $\log_2$ fold change of TSS isoform expression (TSS, ciTSS, aTSS) from full-length mature RNA (A) and nascent RNA (B) across all genes. Rows represent chromatin states and columns represent TSS isoforms. Asterisks denote significance levels (* $p < 0.05$, ** $p < 0.01$, *** $p < 0.001$).

(C) Aggregate nucleosome occupancy profile centered at the +1 nucleosome dyad for high-expression genes with down-regulated TSS isoforms in full-length mature RNA (n = 322 genes), shown for all four samples.

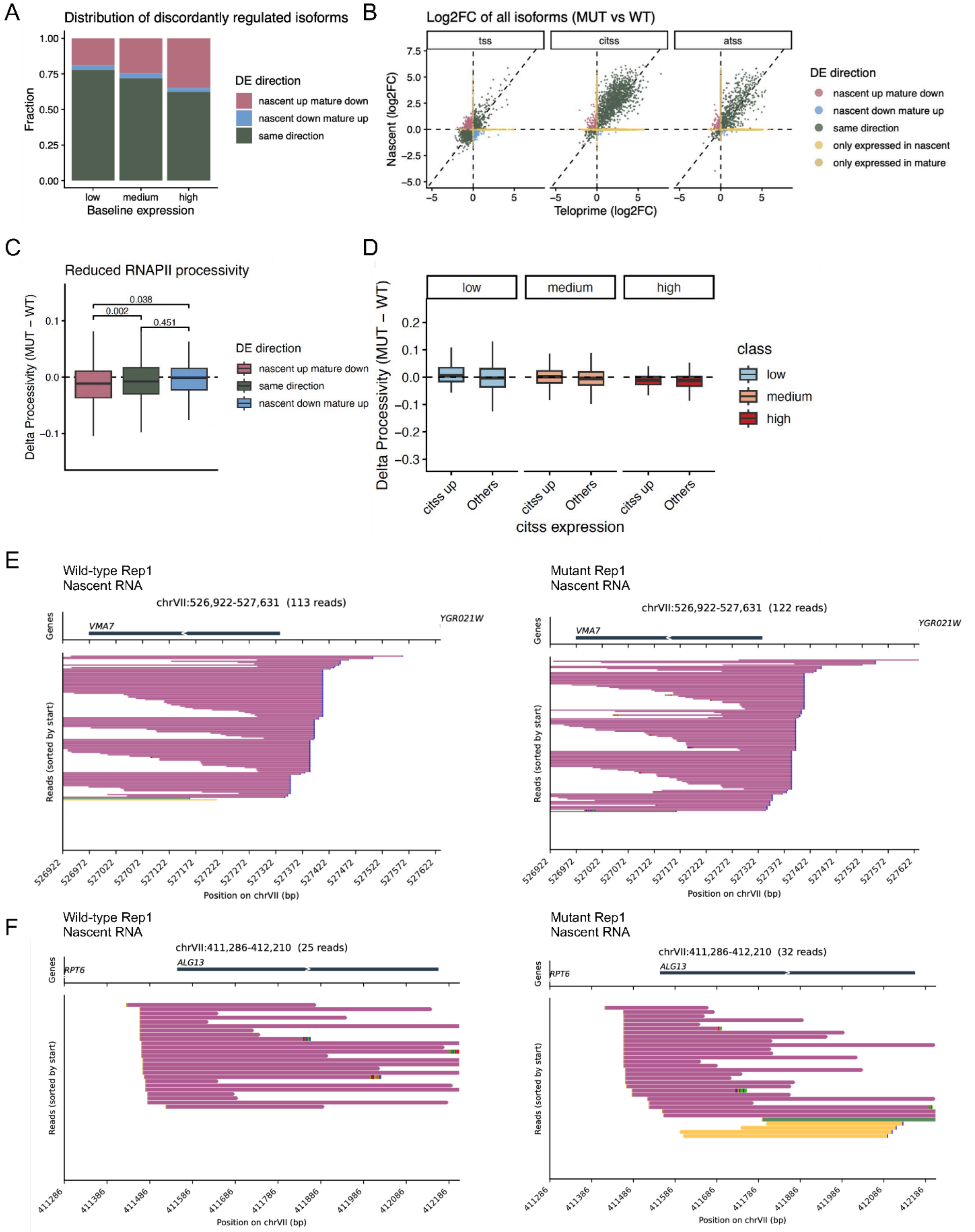


**Supplementary Figure 10. RNAPII processivity defects in *isw1*Δ *chd1*Δ mutant cells.**

(A) Stacked bar plots showing the fraction of TSS isoforms with concordant or discordant regulation between nascent and full-length mature RNA, stratified by baseline expression class

(low, medium, high). The fraction of discordantly regulated genes (nascent up but mature down) is highest in highly expressed genes.
(B) Scatter plots of $log_2$fold change (mutant vs wild-type) in full-length mature RNA versus nascent RNA, shown separately for TSS, ciTSS, and aTSS isoforms. Each point represents one isoform colored by concordance of differential expression.
(C) Boxplots of RNAPII processivity index change (Δ processivity = mutant - wild-type) stratified by concordance of differential expression between nascent and mature RNA. Isoforms discordantly up-regulated in nascent but down-regulated in mature RNA show the greatest processivity loss (Wilcoxon p = 0.002 and 0.038).
(D) Boxplots of delta processivity comparing genes with up-regulated ciTSS expression versus all other genes, stratified by baseline expression class. No significant difference is observed, indicating that processivity loss is not primarily driven by collision with ciTSS-initiating polymerases.
(E-F) Representative genome browser track views of nascent RNA reads at the *VMA7* (E) and *ALG13* (F) loci in wild-type (left) and *isw1*Δ *chd1*Δ mutant (right) replicate1 sample. Progressive coverage loss toward the 3' end is visible in the mutant at individual loci.

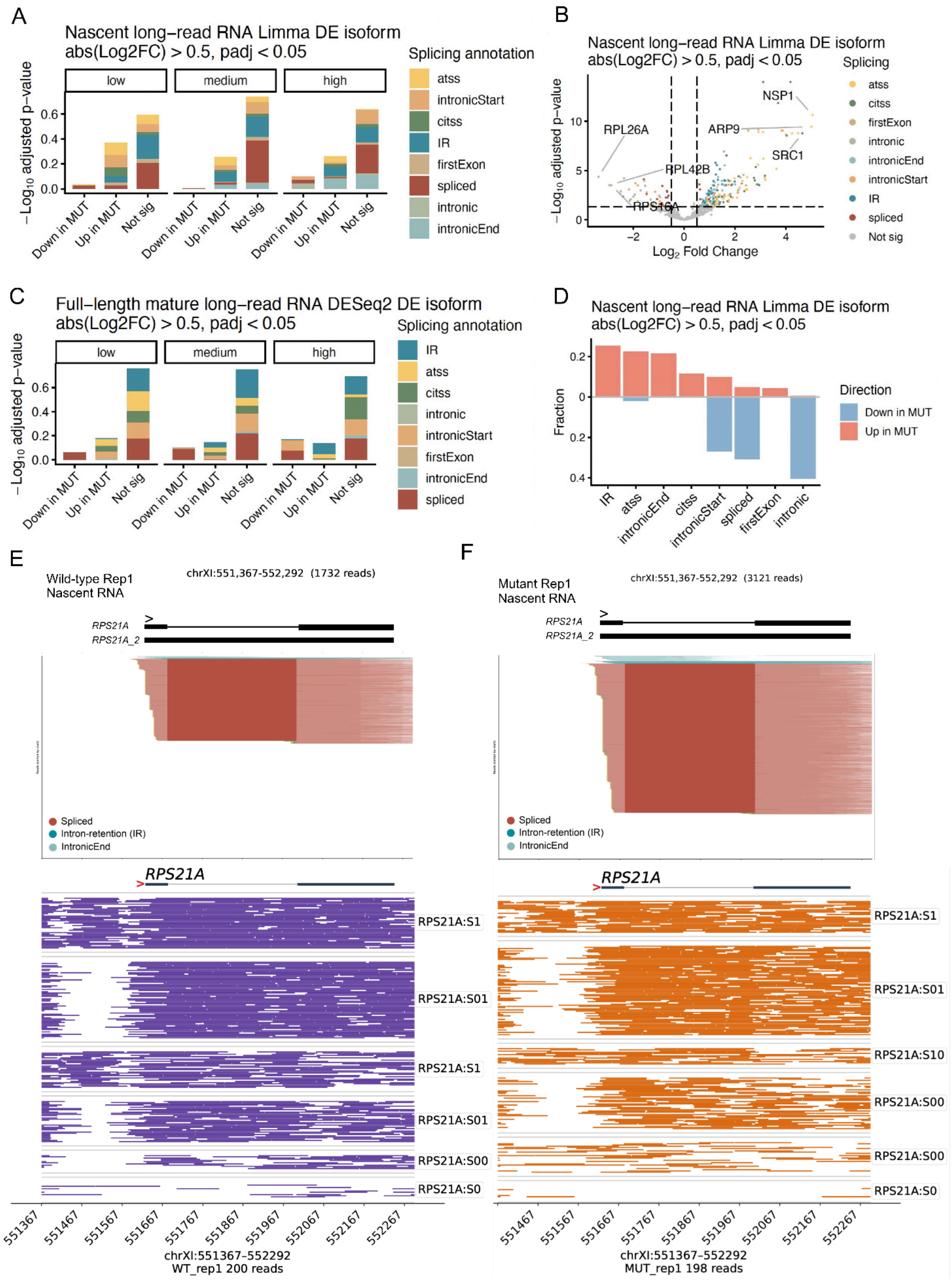

A
Nascent long−read RNA Limma DE isoform
abs(Log2FC) > 0.5, padj < 0.05
low
medium
high
−Log10 adjusted p−value
Down in MUT
Up in MUT
Not sig
Splicing annotation
atss
intronicStart
citss
IR
firstExon
spliced
intronic
intronicEnd
B
Nascent long−read RNA Limma DE isoform
abs(Log2FC) > 0.5, padj < 0.05
NSP1
RPL26A
ARP9
SRC1
RPL42B
RPS16A
Log2 Fold Change
Splicing
Not sig
C
Full−length mature long−read RNA DESeq2 DE isoform
abs(Log2FC) > 0.5, padj < 0.05
D
Nascent long−read RNA Limma DE isoform
abs(Log2FC) > 0.5, padj < 0.05
Fraction
Direction
Down in MUT
Up in MUT
E
Wild-type Rep1
Nascent RNA
chrXI:551,367-552,292 (1732 reads)
RPS21A
RPS21A_2
Spliced
Intron-retention (IR)
IntronicEnd
RPS21A:S1
RPS21A:S01
RPS21A:S00
RPS21A:S0
chrXI:551367-552292
WT_rep1 200 reads
F
Mutant Rep1
Nascent RNA
chrXI:551,367-552,292 (3121 reads)
RPS21A:S10
MUT_rep1 198 reads

**Supplementary Figure 11. Differential splicing isoform usage in nascent and full-length mature RNA.**

(A) Stacked bar plots showing the distribution of differentially expressed splicing isoforms from nascent RNA ($|log_2FC| > 0.5$, $padj < 0.05$, limma), stratified by baseline expression class (low, medium, high) and direction of change, colored by splicing category.

(B) Volcano plot of differentially expressed ($|log_2FC| > 0.5$, $padj < 0.05$, limma) splicing isoforms from nascent RNA. Each point represents one isoform colored by splicing annotation.

(C) Stacked bar plots showing the distribution of differentially expressed splicing isoforms from full-length mature RNA (|log2FC| > 0.5, padj < 0.05, DESeq2), stratified by expression class and direction of change.

(D) Bar plot showing the fraction of up- and down-regulated isoforms by splicing category in nascent RNA.

(E-F) Genome browser track views of nascent RNA reads (top) and single-molecule chromatin fiber maps (bottom) at the *RPS21A* locus in wild-type rep1 (E) and *isw1*Δ *chd1*Δ mutant rep1 (F). Reads are colored by splicing isoform class and sorted by start position. Chromatin fibers are grouped by chromatin state. In the mutant, the intronicEnd isoform increases and chromatin shifts toward intragenic-accessible (S10) and co-accessible (S00) states, illustrating the coupling between chromatin state transition and splicing impairment at single-molecule resolution.

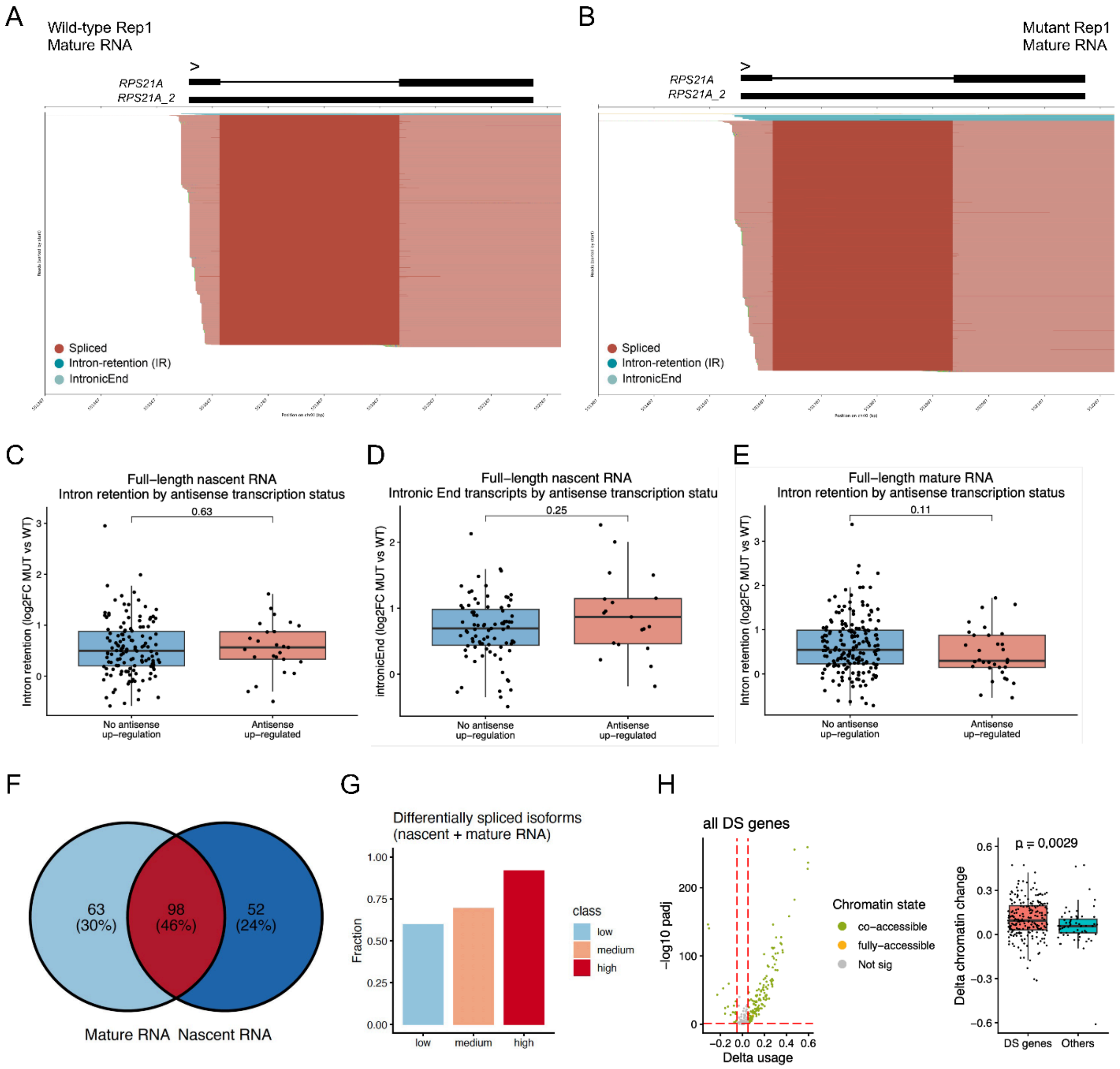


**Supplementary Figure 12. Differential splicing isoform usage and its correlation with aTSS expression and chromatin state alterations**

(A-B) Genome browser track views of full-length mature RNA reads at the *RPS21A* locus in wild-type (A) and *isw1*Δ *chd1*Δ mutant replicate1 sample (B). Reads are colored by splicing isoform annotation and sorted by start position.

(C-D) Boxplot comparing $\log_2$fold change (mutant vs wild-type) of intron retention (C) and intronicEnd (D) isoforms in nascent RNA between genes with and without up-regulated aTSS isoform (Wilcoxon $p = 0.63$ and $p = 0.25$), showing no significant association between up-regulation of antisense transcription and intron retention.

(E) Boxplot comparing $\log_2$fold change (mutant vs wild-type) of intron retention isoform in mature RNA between genes with and without up-regulated aTSS isoform (Wilcoxon $p = 0.11$), showing no significant association between up-regulation of antisense transcription and intron retention.

(F) Venn diagram showing overlap between genes with differentially spliced isoforms in full-length mature RNA (n = 161) and nascent RNA (n = 150). Of 213 total differentially spliced genes, 98 (46%) are detected in both libraries.
(G) Bar plot showing the fraction of differentially spliced genes in each expression class (low, medium, high). Differentially spliced genes are enriched in highly expressed genes.
(H) Left: volcano plot showing differentially enriched chromatin states (picked one state per gene based on the smallest p-value) for genes with differential splicing (DS genes). Right: boxplot comparing the delta chromatin state (picked one state per gene based on the smallest p-value) change between DS genes and all other genes. DS genes show significantly greater chromatin state gain than other genes (Wilcoxon p = 0.0029).
